# NucLogic: Logic-based geometric 3D nuclear segmentation for fast and robust analysis of challenging biological imaging data

**DOI:** 10.64898/2026.10.05.756154

**Authors:** Sebastian G. van Dijk, Mario Ledesma-Terrón, Susanne J. Kraus, Ruth van Brussel, Emmanuel Marquez-Legorreta, Lars J.S. Kemp, Saskia J.E. Suijkerbuijk

## Abstract

Nuclear segmentation in 3D microscopy remains challenging, particularly for low signal-to-noise ratio (SNR) data from dense tissue samples, timelapse imaging, or in vivo experiments. Yet accurate 3D nuclear segmentation is a prerequisite for quantitative single-cell analysis, making this a key bottleneck for high-throughput study of challenging datasets. We present NucLogic, a fast and robust 3D nuclear segmentation algorithm that combines the maturity of 2D segmentation with geometric stitching rules based on nuclear shape, orientation, and intensity, to reconstruct accurate 3D volumes. NucLogic reliably segments challenging datasets including long-term timelapse imaging of organoids, and large dense structures such as the zebrafish brain. Benchmarked against other state-of-the-art segmentation tools, NucLogic matches F1 scores at high SNR, with increasing outperformance as SNR decreases. Importantly, segmentation completes approximately 3-to 5-fold faster, with greater speed gains on larger images. Beyond segmentation, NucLogic includes an intuitive graphical user interface allowing non-experts to segment, quantify, and visualize 3D and 4D data. This enables straightforward quantification of cell numbers, alongside integration of a wide range of both cell-intrinsic properties, such as phenotype and morphology, and local environment metrics, such as density and spatial interactions. Together, NucLogic makes quantitative 3D nuclear analysis of previously unmanageable datasets accessible to a broad research community.

## Introduction

Understanding how tissues form, function, and respond to perturbation requires the ability to resolve and quantify individual cells or nuclei within their native, densely packed three-dimensional context. Robust and accurate nuclear segmentation is needed to study these spatiotemporal mechanisms. Recent advances in deep-learning-based approaches, such as Cellpose [1,2], StarDist [3,4], and PlantSeg [5], have enabled automated segmentation of biological images. Especially on 2D images, modern AI-based algorithms often outperform humans and conventional image analysis methods, in both speed and accuracy. However, generating accurate 3D segmentations has remained challenging due to increased sample complexity, reduced imaging quality of thick samples, and the extensive manual effort required to generate annotated 3D training data. This challenge is especially pronounced in highly dynamic tissues and three-dimensional culture systems, such as organoids, whose imaging is often characterized by low signal-to-noise ratio (SNR), limited z-resolution, and z-dependent intensity loss. Timelapse imaging poses an additional challenge, as minimizing phototoxicity and total acquisition time typically requires trading off image quality for speed [6]. Because of this, even after model fine-tuning, existing deep-learning-based 3D approaches fail to accurately segment images with poor SNR resulting from a drop in signal intensity along the z-axis, as is common in confocal imaging. Moreover, most current segmentation and downstream image analysis tools remain difficult to configure and operate for non-experts, and the resulting segmentation process itself is often both time- and compute-intensive. Combined, this has restricted the broader adoption of quantitative 3D image analysis across the life sciences [7].

To address these limitations, we developed NucLogic, which leverages the robustness of existing 2D segmentation approaches with a novel stitching algorithm to achieve accurate nuclear segmentation. The idea of a slice-by-slice approach to 3D segmentation has previously been described, however, each strategy comes with crucial limitations. For example, Cellpose [2] creates 3D segmentation by segmenting along all XY, YZ and XZ planes. Besides being computationally expensive, this leads to suboptimal results, as XZ and YZ planes suffer from a low z-resolution (4 to 10-fold lower compared to XY) and segmentation along the YZ and XZ planes is impaired by decreasing brightness along the z-axis. u-Segment3D [8] follows a similar 2D-to-3D strategy, but, in our benchmarking, does not reliably segment large volumes of densely packed nuclei. Alternatively, OSCAR [9] stitches existing 2D XY segmentations into 3D clusters by linking PCA-derived 2D projections across adjacent z-slices with a feature-vector machine learning algorithm, using statistical rules that let a single 2D object contribute to multiple clusters to tolerate low z-resolution and plane-by-plane errors. However, its parametric 3D fits are designed to determine object position and orientation, rather than to capture actual nuclear morphology.

In contrast, NucLogic applies Cellpose-SAM exclusively in the XY plane, mitigating the effects of anisotropic resolution and z-dependent intensity variation, and then transforms the resulting 2D masks into 3D volumes that retain individual nuclear morphology using a set of human-interpretable geometric rules. These rules are based on cross-slice area, intensity, and orientation patterns, offering a transparent alternative to the black-box nature of most deep-learning-based 3D segmentation approaches. This approach substantially outperforms existing methods in both accuracy and speed on complex samples. NucLogic achieved remarkably high recall in low-SNR areas of 3D image stacks on both synthetic and experimental data. Moreover, segmentation is performed approximately 3-fold faster than existing methods, making NucLogic particularly well suited to time-sensitive applications, such as high-throughput screening or smart microscopy [10].

To enable broader adoption of 3D segmentation analysis across the biology field, NucLogic is designed to be accessible to non-expert users, making accurate nuclear segmentation accessible to a broader range of researchers and advancing the field as a whole. To this end, the NucLogic suite includes an easy to navigate graphical user interface (GUI) that supports data cropping, visualization, and extraction of a range of statistics, without requiring computational experience. The NucLogic GUI enables extensive data mining of complex datasets at single-cell resolution to profile both cell-intrinsic and spatial phenotypes. This allowed us to identify and quantify growth rate heterogeneity across organoids, driven by differences in spatial organization and cell-cell interactions. Our results demonstrate that NucLogic has the potential to make reproducible, single-cell-resolution analysis of complex, low-SNR, multi-dimensional imaging data broadly accessible to biologists, facilitating the discovery of novel spatiotemporal patterns across large-scale datasets.

## Results

### A geometric ruleset accurately constructs 3D nuclear volumes from 2D masks

Despite recent advances in deep-learning approaches, 3D segmentation remains challenging. Even fine-tuned 3D segmentation models often fail to accurately segment images with poor SNR and z-dependent intensity loss. Intriguingly, we noticed that the individual slices of such image stacks are correctly segmented by 2D segmentation models (Fig. 1A). Therefore, we hypothesized that a slice-by-slice segmentation approach using existing 2D methods with a specialized stitching algorithm would enable accurate segmentation of these challenging stacks.

**Figure 1:**
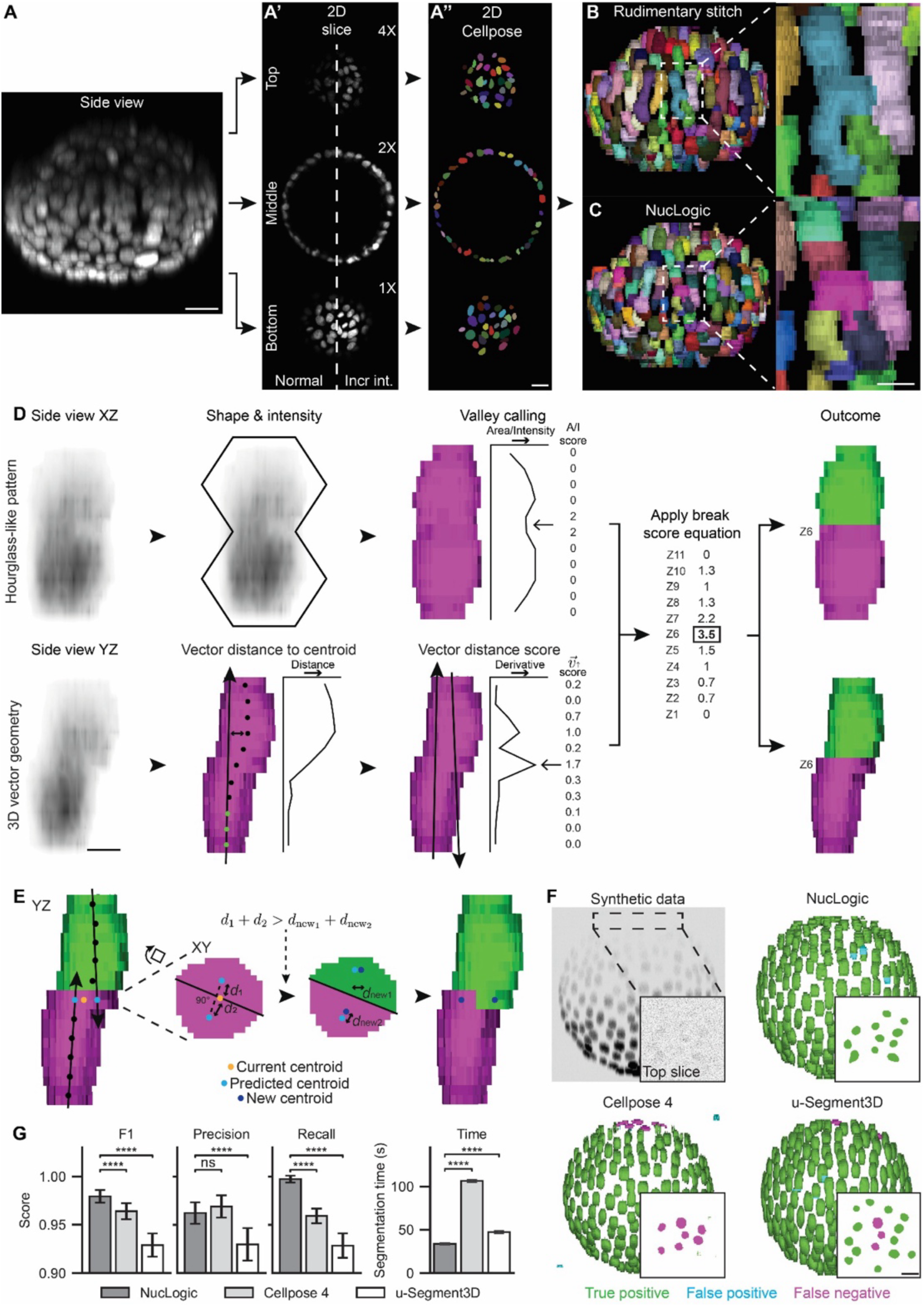
A geometric ruleset accurately constructs 3D nuclear volumes from 2D masks. (A) Left image shows a sideview of a 3D reconstructed colorectal cancer organoid. Nuclei (gray) are visualized with H2B-mCherry. A’ displays individual z-planes of the bottom, middle and top of the z-stack with similar (left) or 1x/2x/4x increased contrast settings. A’’ shows its corresponding 2D segmentation masks generated with Cellpose-SAM. (B) The 2D Cellpose-SAM masks can be stitched together using a rudimentary IoU stitch approach. Zoomed image displays z-undersegmented nuclei. (C) NucLogic is applied to break the undersegmented nuclei into correct 3D objects. Scale bar = 5 µm. (D) Geometric criteria for identifying break points in z-undersegmented nuclei. Top panel shows an XZ sideview of two z-undersegmented nuclei. The area and intensity constriction follows an hourglass-like pattern that can be used to identify the optimal break point by identifying valleys in the segmented mask in pink. Bottom panel shows the same two nuclei from YZ perspective. The 3D orientation vector is fitted to the three bottom most centroids to calculate distance and the consecutive vector distance score. The right image displays how these scores are combined into a final per-z-slice break score to identify the optimal break point and the corresponding two segmented nuclei. Scale bar = 5µm. (E) On the left a YZ sideview shows the same segmented nuclei. Inward-facing 3D vectors predict the expected centroid position in the touching slice. A split is accepted when it reduces the distance between predicted and resulting centroids as shown in the center images. On the right, shared 2D segmentations at the interface between touching nuclei are redistributed to their correct objects. (F) Side view of a representative synthetic organoid (top left) and segmentation masks generated by NucLogic, Cellpose-SAM, u-Segment3D. True positive shown in green, false positives shown in blue, false negatives shown in magenta, insets show a zoom of the top slice. Scale bar = 25 µm. (G) Quantification of the F1, precision, and recall score, as well as the segmentation time per synthetic organoid (N = 20) across all three benchmarked methods. Data are presented as mean ± SD.

We selected Cellpose-SAM (also known as Cellpose-4) for 2D segmentation, as this model showed the most robust segmentation on low-SNR slices (Fig. S1). Basic algorithms exist to transform a stack of 2D segmented slices into 3D objects via rudimentary stitching approaches (Fig. 1A’-B). Most of these rely on the Intersection over Union (IoU) between 2D masks in neighboring slices, defined as the ratio between the overlapping area and the total combined area of the two 2D masks. However, applying this to dense 3D tissue stacks leads to undersegmentation along the z-dimension (Fig. 1B). This is a direct consequence of the limited axial resolution of a confocal microscope, caused in part by the elongation of the point-spread function (PSF) along the z-axis, penetration depth and light scattering through a volumetric sample (Fig. 1A). To overcome this, we set out to define a human-interpretable set of rules that can reliably transform stacks of 2D nuclear segmentation masks into 3D volumes, even in densely packed tissues (Fig. 1C-D).

At its core, NucLogic splits densely z-undersegmented nuclei into individual objects by assessing each initial nuclear mask against two geometric and intensity-based criteria, which determine whether a split is required and, if so, at which slice it should occur. First, objects containing z-undersegmented nuclei often follow an hourglass-like pattern, with the optimal breaking point found at a local minimum in both 2D slice area and relative intensity (Fig. 1D). Local minima are identified by comparing the area and intensity of each 2D mask to its neighboring slice masks, assigning a break-likelihood score to each slice (termed *A* for area and *I* for intensity). The break score reflects the relative constriction in area and intensity observed between z-stacked nuclei. Intensity break scores are relative to the intensity at its minimum, which makes the score invariant to signal heterogeneity throughout the z-axis.

Secondly, individual nuclei in a z-undersegmented object often have slight variations in 3D orientation. To assign a mathematical description to this, NucLogic estimates the orientation of the lower nucleus by fitting a 3D vector to the centroids of the three bottom-most slices of a z-undersegmented object using least-squared regression (Fig. 1D). The distance between each 2D centroid and the vector describes the likelihood that the corresponding slice belongs to the lower nucleus [9]. We extend this approach by using the rate of change of this distance (i.e., its derivative) to convert this geometric measure into a vector-based break-likelihood score (*v*^⤍^↑). The slice at which this distance most rapidly increases (i.e. the maximum of the derivative) reflects the appropriate breaking point between two z-undersegmented nuclei. Calculating this vector-based score in both the upward and downward orientation enables finding the most accurate break point, even when three or more nuclei are stacked. Next, all individual scores are combined into a final per-slice break score (*S_break_*) using *Equation 1* (Fig. 1D). Additionally, an altered version of the Intersection over Union (*IoU*) of each slice with its top neighbor is incorporated in *S_break_* to penalize stacked nuclei with similar orientation but a slight misalignment in XY (see methods). Stacked nuclei containing a slice with an *S_break_* above a set breaking threshold (*T_break_*) are split between the two slices with the highest *S_break_*(Fig. 1D). The applied *T_break_* is calculated individually for each nucleus based on its elongation (*Equation 2*), further promoting the splitting of z-stacked nuclei. Moreover, the initial breaking threshold can be refined on a per-tissue-type basis to account for differences in z-density and axial blur. However, this is generally unnecessary as a wide range of initial thresholds yield similar segmentation outcomes due to the elongation-based recalculation of *T_break_*. All nuclei in the sample are broken iteratively until no more nuclei contain slices with *S_break_* > *T_break_*. A final cleanup step removes small background artifacts, defined as single- or dual-slice objects that cannot be reasonably attributed to a nucleus.

As this slice-by-slice approach is dependent on 2D segmentation, any 2D undersegmentation errors are directly translated into 3D segmentation errors. This leads to the attribution of a single undersegmented slice to one nucleus instead of correctly subdividing it over the two z-stacked nuclei (Fig. 1E). To resolve this, we implemented a method to split shared 2D segmentations. For each pair of touching nuclei, inward-facing 3D vectors are generated from the 2D centroids of all slices except the touching slice. These vectors are used to predict the expected centroid location within the touching slice. Next, we compare two distances in the touching slice: (1, *dpred*) the distance between the current (unsplit) centroid and the predicted centroids, and (2, *dnew*) the distance between the predicted centroids and the centroids that would result from splitting the touching slice perpendicular to the predicted centroids (Equation 3). When the potential split leads to a decreased distance to the predicted centroids, the split is accepted (Fig. 1E). After a successful split, the next consecutive z-layer is also checked for undersegmentation in a similar manner. Combined, these geometric criteria enable reliable reconstruction of any stack of 2D nuclear segmentations into individual 3D nuclei.

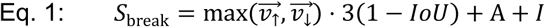

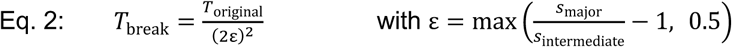

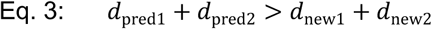

### NucLogic outperforms existing segmentation methods on synthetic organoids under z-dependent SNR loss

Next, we generated in-silico data to quantitatively assess the performance of NucLogic’s 3D segmentation. Synthetic nuclear-labeled organoids were generated by randomly distributing ellipsoidal nuclei across the surface of a sphere, each oriented towards the organoid center to mimic the in-vitro characteristics of cystic organoids. Depth-dependent optical blurring, signal attenuation, and Poisson-Gaussian noise were then applied to each image stack to mimic the z-axis resolution loss and SNR degradation observed in confocal images (see Methods). Generated organoids had a diameter of approximately 160 µm and contained nuclei averaging 6×15×15 µm. Volumes were sampled at an anisotropic voxel size of 2.5×0.64×0.64 µm (ZYX) to reflect typical confocal acquisition settings.

To benchmark NucLogic against existing state-of-the-art algorithms, we segmented 20 synthetic organoids using NucLogic, Cellpose-SAM [1], u-Segment3D [8], and StarDist [3]. Cellpose-SAM was run using its built-in 3D mode configured with the same nuclear-pretrained model used for NucLogic. u-Segment3D was selected for benchmarking as it stitches 2D masks of Cellpose into 3D segmentations, similar to NucLogic’s approach. However, u-Segment3D relies on 3D gradient vectors, combined with gradient descent and spatial connected component analysis to generate 3D segmentations, rather than the human-interpretable geometric rules used by NucLogic [8]. For segmentation, u-Segment3D was applied in an indirect manner, supplying our own 2D Cellpose-SAM masks rather than using its native pipeline, as this produced better results than the direct u-Segment3D framework when configured with either Cellpose 3 or Cellpose-SAM models. StarDist was performed using the pretrained 3D model, as training a custom model is not yet well supported.

Across all 20 synthetic organoids, NucLogic achieved the highest segmentation accuracy of the four tested methods (Fig. 1F-G). Only a small number of segmented false-positive nuclei were observed, primarily resulting from over-splitting at the sides of the organoid (Fig. 1F). Resulting in a near-perfect precision score of 0.962 ± 0.011 (Fig. 1G and Table S1). Notably, almost no false negatives were observed, and an almost perfect recall score was achieved (0.997 ± 0.004, mean ± SD), leading to the highest overall F1 score (0.979 ± 0.007, Fig. 1G). In contrast, Cellpose-SAM 3D showed a mixture of both false-negative and false-positive nuclei, resulting in a slightly lower F1 score (0.964 ± 0.008, Fig. 1G). As expected, false-negative nuclei were mainly present at the top of the z-stack, where nuclear intensity was relatively weak. u-Segment3D exhibited a less predictable pattern of false-negative and false-positive cells (F1 = 0.929 ± 0.012, Fig. 1G). StarDist performed particularly poor on this low-SNR image with an average F1 score of 0.568 ± 0.019 (Fig. S2, Table S1).

Additionally, NucLogic was the fastest well-performing segmentation algorithm, with an average segmentation time of 34 seconds per synthetic organoid (Fig. 1G). This corresponds to a 3-fold speed improvement over Cellpose-SAM (106 seconds) and a 1.5-fold improvement over u-Segment3D (47 seconds). Taken together, these results demonstrate that NucLogic outperforms existing state-of-the-art methods in both accuracy and speed when segmenting nuclei in synthetic low-SNR confocal images.

### NucLogic accurately segments various tissue types at varying SNR levels

To further benchmark NucLogic, we turned to experimental NIS3D dataset, containing fully annotated masks of the Drosophila and zebrafish embryo [11]. Segmentation of embryonic nuclei is exceptionally challenging, as these samples are characterized by high nuclear density, large imaging volumes, low SNR, and a variety of nuclear shapes and intensities. The annotated Drosophila embryo is a relatively high-quality image stack, featuring clearly separated nuclei and consistent image quality throughout the z-stack. In this scenario, both NucLogic and Cellpose-SAM show highly accurate segmentation with insignificant differences in F1 score (Fig. 2A, Table S2). Additionally, errors at this level of performance are partially explained by interpretation differences and annotation mistakes by human annotators [11]. Both u-Segment3D and StarDist underperformed relative to NucLogic, even on these high-quality samples. Notably, NucLogic completed segmentation ∼2.5-fold faster than Cellpose-SAM, recapitulating the speed advantage observed in the synthetic benchmark (Fig. 2A).

**Figure 2:**
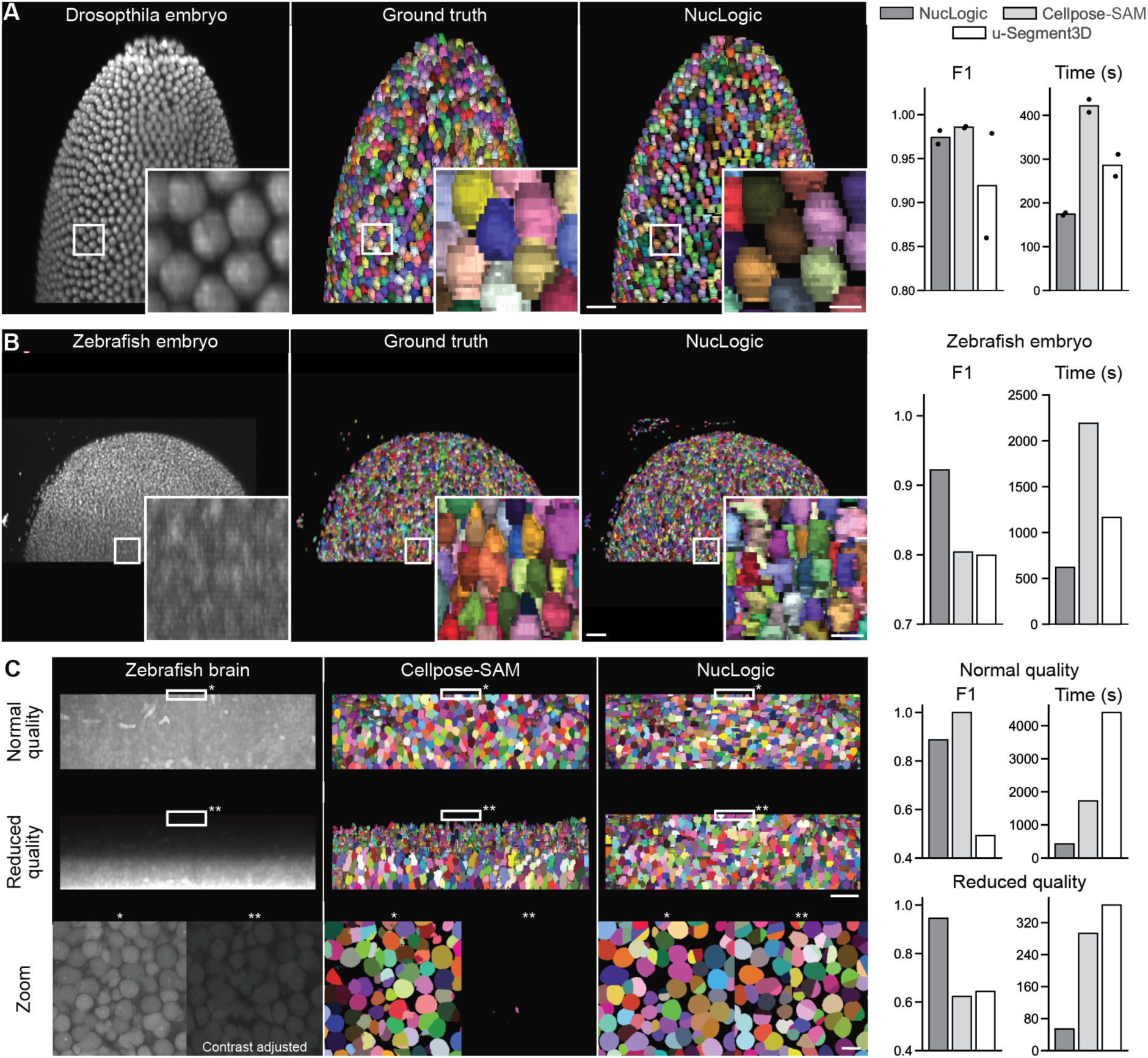
NucLogic accurately segments various tissue types at varying SNR levels. (A) Left image shows a representative Drosophila embryo from the NIS3D dataset. Middle image shows ground truth segmentation as derived from NIS3D. Right image shows corresponding NucLogic segmentation. Right graph displays segmentation accuracy and speed for NucLogic, Cellpose-SAM, u-Segment3D using the centroid accuracy method (see methods), each dot represents an embryo. (B) Similar representation of the zebrafish embryo of the NIS3D dataset. (C) Left images show the expansion-microscopy-imaged zebrafish brain volume at normal and reduced quality, alongside zooms of the top-most z-slices of the data. Middle images show Cellpose-SAM segmentation of these samples. Right images show NucLogic segmentations. Graphs display F1 segmentation accuracy (centroid accuracy) and segmentation time of NucLogic, Cellpose-SAM, and u-Segment3D for the normal and reduced quality images. F1 of Cellpose-SAM at normal quality is set to 1 because it was considered the ground truth segmentation. Scale bars are 50 µm for full images and 10 µm in zoomed images.

The zebrafish embryo of the NIS3D dataset presents a more challenging, low-SNR stack with variable intensity along the z-axis. Consistent with the synthetic benchmark, NucLogic achieved a significantly higher F1 score than existing methods on this more demanding sample type (Fig. 2B). This can largely be attributed to NucLogic’s high recall (0.945 compared to 0.689 for Cellpose-SAM, Table S3). Cellpose-SAM failed to detect a substantial proportion of nuclei in the deeper, lower-intensity regions of the embryo, whereas NucLogic successfully identified most of these nuclei (Fig. S3). We additionally observed that NucLogic-generated segmentations more closely matched the shape of individual ground-truth nuclei, confirmed by consistently higher F1 scores at increasing IoU thresholds when using Hungarian-algorithm-based matching (Fig. S4). Of note, NucLogic detected substantially more correct nuclei than Cellpose-SAM (13,414 nuclei with an F1 of 0.922, versus 9,130 nuclei with an F1 of 0.804) and also completed segmentation approximately 3.5-fold faster.

Next, we aimed to rigorously evaluate NucLogic’s computational capabilities by segmenting an ultra-dense and large-volume sample. To this end, we performed expansion confocal microscopy on the zebrafish brain. The dataset has a volume size of 101×2048×2048 pixels and a voxel size of 1×0.16×0.16 μm (not including 2x expansion), containing approximately 14,000 nuclei (Fig. 2C). A high SNR is achieved throughout the sample by optical clearing of the tissue during expansion, resulting in a very stable intensity throughout the z-axis. Because of this, Cellpose-SAM achieved near-perfect segmentation, and we considered this the ground truth segmentation. At this resolution, some of the geometric rules that form the basis of the NucLogic algorithm fail to accurately represent nuclear shapes. Likely, nuclear shape is too irregular at this resolution, generating an hourglass-like pattern within a single nucleus instead of between z-stacked nuclei. This resulted in a significant drop in segmentation accuracy of NucLogic compared to Cellpose-SAM (Fig. 2C, Table S4). Nevertheless, NucLogic segmented this challenging volume in considerably less time than the other methods (∼4 and ∼10-fold faster than Cellpose-Sam and u-Segment3D, respectively). This demonstrates that NucLogic’s underlying algorithm remains robust and scales favorably with large image volumes.

The availability of near-ground-truth masks for this dense tissue presented a unique opportunity to investigate segmentation of ultra-dense tissues under relatively poor image quality such as that acquired during timelapse imaging. To this end, we manipulated the expanded zebrafish brain sample to emulate real-world confocal imaging conditions using the same approach previously applied to generate synthetic data. In addition, the volume was down sampled two-fold along each axis (voxel size of 2×0.32×0.32 μm). This allowed us to approximate the imaging conditions typical of conventional, non-expanded confocal microscopy of similar tissue. Upon image quality reduction, NucLogic again compared favorably to the other tested methods and accurately segmented nuclei in low-intensity slices, whereas other methods did not (Fig. 2C, Table S5). Collectively, these findings demonstrate that NucLogic is exceptionally well-suited for nuclear segmentation in an assortment of challenging dense image stacks, surpassing existing state-of-the-art algorithms in both accuracy and speed.

### A GUI and general pipeline for image analysis

Beyond addressing the segmentation challenges, NucLogic is designed to be broadly accessible to non-coding users, making accurate nuclear segmentation accessible to a broader range of researchers and advancing the field as a whole. Therefore, the NucLogic suite includes an easy to navigate graphical user interface (GUI) that supports data cropping, visualization, and extraction of a range of statistics, without requiring computational experience (Figs. 3 and S5). NucLogic supports all major image file formats, accommodating single- or multi-timepoint acquisitions with one or more channels. Processing begins with cropping of the input data, performed automatically for timelapse data, and either automatically or manually for static imaging data (Figs. 3 and S6). Restricting the cropped region to a single structure of interest, with minimal surrounding background ensures that downstream statistics are calculated only for nuclei belonging to that structure, rather than being pooled across multiple structures within the same field of view. This also reduces computational load and downstream data storage requirements. Automatic cropping of timelapse data is performed in the XY dimensions by using a locally run implementation of Meta’s SAM 2 video segmentation model, which tracks large individual structures, such as organoids, across timelapses containing multiple structures within the field of view [12]. Cropping along Z is performed in a slice-by-slice manner by calculating a Moran’s I score for spatial autocorrelation to distinguish foreground slices that contain non-random signal from background slices containing random noise. Alternatively, XY cropping can be performed manually by drawing a region of interest on the z-projection of the sample.

**Figure 3:**
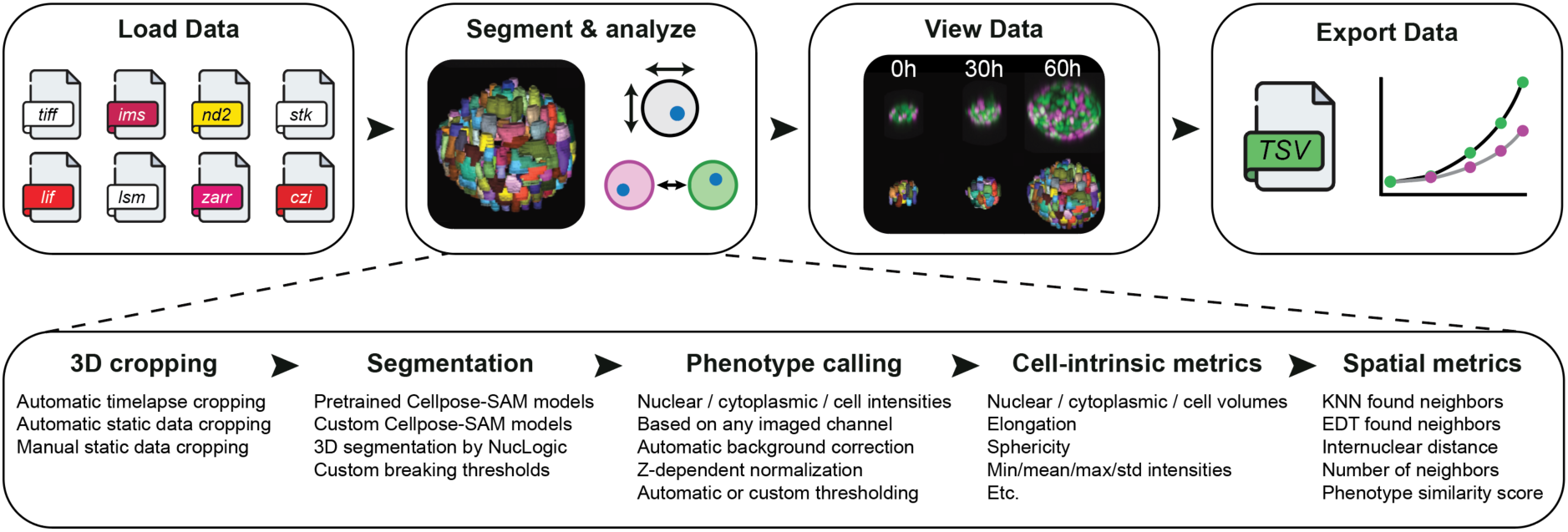
A GUI and general pipeline for image analysis. The top panel depicts the overall flow of the graphical user interface (GUI) of NucLogic. The GUI comprises four main tabs for loading, segmenting, viewing, and exporting data. Multiple input image formats are supported. Data can be visualized using napari. Full or summarized data can be exported as TSV files for further processing and plotting. The bottom panel shows the range of features available through the NucLogic UI. 3D cropping, segmentation, phenotype calling, and the acquisition of numerous cell-intrinsic and spatial metrics can all be configured and performed through the GUI.

Following cropping, the sample is segmented using 2D Cellpose-SAM in combination with the NucLogic stitching algorithm. Subsequently, NucLogic can apply a built-in phenotype-calling algorithm that assigns a phenotype to each segmented nuclei based on any imaged fluorescent marker, enabling straightforward quantification of user-defined phenotypes of interest. Moreover, NucLogic provides a wide range of statistics, including nucleus-intrinsic properties including size, location, morphology, and intensity, as well as spatial context-dependent statistics such as cell density and interactions between neighboring cells (Figs. 3 and S5). Finally, the resulting segmentations and their accompanying statistics can be directly visualized by launching napari from the NucLogic UI (Fig. S7). Together, these components form an accessible, end-to-end pipeline that enables accurate nuclear segmentation directly from raw imaging data, even under challenging low-SNR conditions.

### NucLogic enables extensive data mining of cell-intrinsic and local environment metrics at single-cell resolution

To assess the real-world analytical capabilities of the NucLogic suite, we reanalyzed our previously published organoid timelapse dataset [13]. It comprises mixed murine organoids, generated by combining wild-type (WT) liver progenitor organoids with colorectal cancer organoids. Both populations contain the nuclear marker H2B-mCerulean and can be separated by membrane-Tomato (mTmG) and Dendra2 (D2) expression of the WT and cancer cells, respectively (Fig. 4A). Leveraging its integrated cropping, segmentation, and phenotype calling workflow, NucLogic fully segmented and analyzed the entire dataset, containing 157 organoids of 1020×1020×122 16-bit voxels across 14 timepoints each, in under 6 hours on a standard workstation. The available phenotype information enabled direct quantification of the cell-type-specific growth rates of each organoid (Fig. 4A). The observed growth rates of this reanalysis are in line with previously reported findings (Fig. 4B) [13].

**Figure 4:**
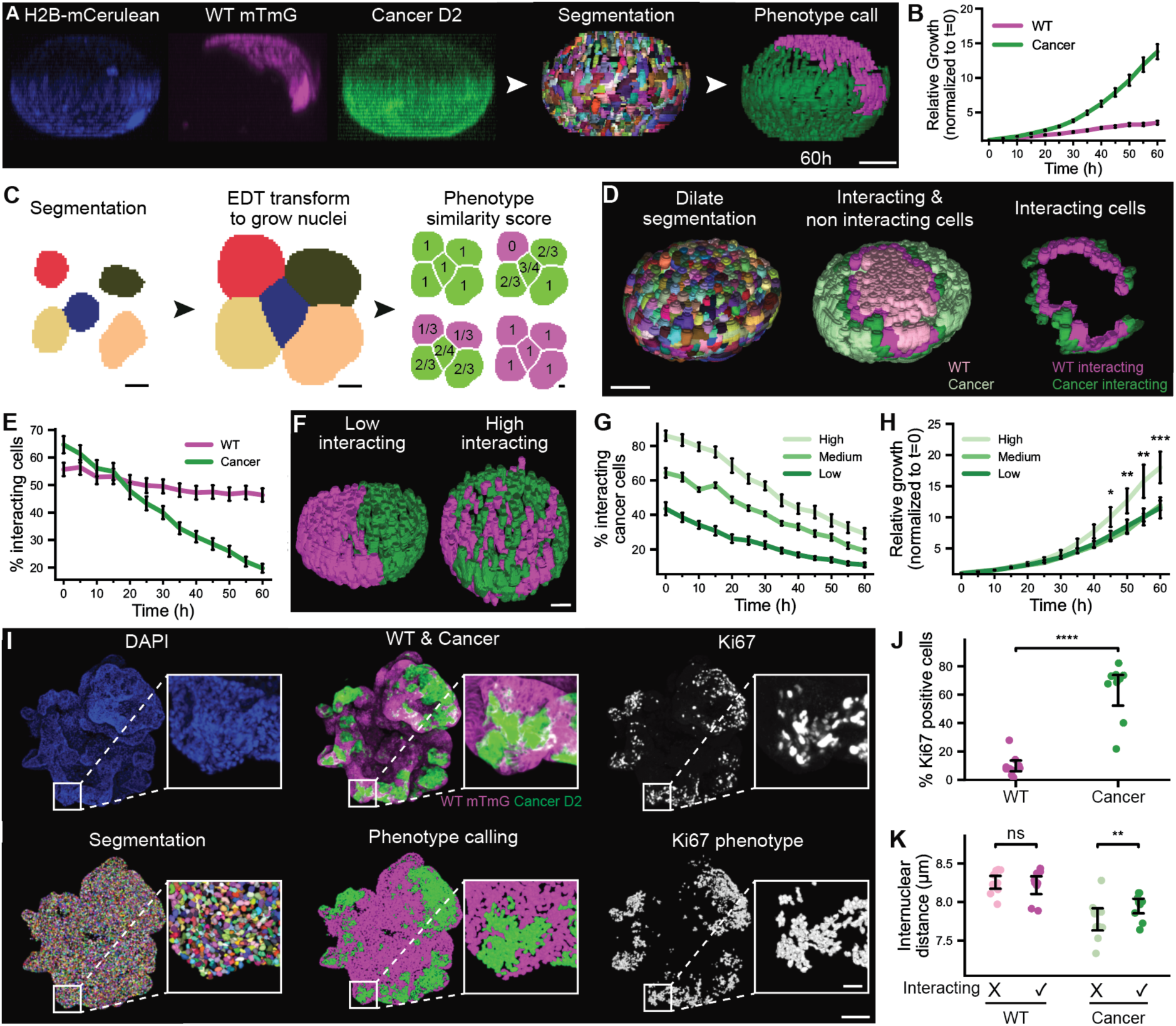
NucLogic enables data mining of cell-intrinsic and local environment metrics at single-cell resolution. A) Mixed murine organoid combining wild-type (WT) liver progenitor organoids labelled with mTmG (magenta) and colorectal cancer organoids labelled with Dendra2 (green). Both populations are labeled with H2B-mCerulean3 (blue). NucLogic-generated segmentation and phenotype calls are shown for each nucleus. Scale bar = 50 µm. (B) Timelapse microscopy dataset of the same organoids analyzed using NucLogic. Population-specific growth rates are quantified via NucLogic’s phenotype-calling workflow and plotted. Data represented as mean ± SEM. N = 46 mixed organoids. (C) Left: example of nuclear segmentation. Middle: EDT-based nuclear mask dilation used to estimate cell-cell contacts. Right: phenotype similarity score (PSS) calculated for representative phenotype combinations. Scale bar = 10 µm. (D) Left: dilated nuclear segmentation of the same organoid shown in panel A. Middle: interacting (PSS < 1) and non-interacting (PSS = 1) cancer and WT cells. Right: interacting cells only. Scale bar = 50 µm. (E) Plot of percentage of interacting cancer and WT cells in mixed organoids over time. Data represented as mean ± SEM. N = 46 mixed organoids. (F) Representative images of mixed organoids with a low (left) and high (right) percentage of interacting cells. Scale bar = 25 µm. (G) Mixed organoids were subdivided into three equal-sized groups based on their mean percentage of interacting cancer cells, and the mean ± SEM percentage of interacting cancer cells per group is plotted (N = 46 mixed organoids). (H) Plot of the normalized growth rate of cancer cells in highly interacting versus low- and medium-interacting organoids. Significance is calculated between high- and low-interacting categories using a linear mixed-effects model. (I) Colorectal liver metastasis microtissues composed of WT (mTmG, magenta) and colorectal cancer (Dendra2, green) cells are stained using DAPI (blue) and Ki67 (grey). NucLogic segmentation, WT versus cancer phenotype calls, and Ki67-positive classification are shown as acquired by NucLogic. Scale bar = 150 µm, 25 µm for zoomed images. (J) Plot of the percentage of Ki67-positive cells in the WT and cancer population (N = 11 microtissues; mean ± SD; independent t-test). (K) Plot of internuclear distance (calculated using three nearest-neighbor density) for WT and cancer cells, separated into interacting and non-interacting populations (N = 11 microtissues; mean ± SD; paired t-test). * = p < 0.05, ** = p < 0.01, *** = p < 0.001, **** = p < 0.0001.

The extensive cell-intrinsic and spatial statistics generated by NucLogic enabled further data mining of this existing dataset. Cell-cell contacts were estimated by emulating whole-cell segmentation, using NucLogic’s built-in 3D Euclidean distance transform (EDT) function to dilate the nuclear masks (Fig. 4C). Combined with its phenotype-calling function, NucLogic calculates a phenotype similarity score (PSS). The PSS represents, on a scale of 0 to 1, the phenotypic similarity of a nucleus to its neighboring cells (Fig. 4C). This metric distinguished interacting cancer cells (PSS < 1) from non-interacting cancer cells (PSS = 1) and enabled the same distinction for WT cells relative to their cancer-cell neighbors (Fig. 4D). Using this analysis, we observed that the percentage of interacting cancer cells within each organoid reduced significantly over time, while the percentage of interacting WT cells remains relatively stable (Fig. 4E). In addition, organoids could be stratified into those containing a low or high percentage of interacting cells, reflecting variability in the irregularity of the interaction front (Fig. 4F). This organoid-level characteristic remained stable over time, rather than reflecting a time-dependent process (Fig. 4G). Intriguingly, cancer cells in highly interacting mixed organoids showed an increased growth rate compared to those in low- or medium-interacting organoids (Fig. 4H; 18.0 ± 1.7 versus 11.5 ± 2.5 fold relative growth at 60h for highly and lowly interacting organoids, mean ± SEM, p = 0.0003). This suggests that the increased proliferative capacity of mixed cancer cells is primarily driven by short-ranged cell-cell interactions with WT cells. Such findings illustrate how NucLogic enables the discovery of novel spatial patterns in cellular behavior that would otherwise remain inaccessible, owing to its combination of robust 3D timelapse segmentation and friendly workflow.

Next, we assessed NucLogic’s ability to analyze data of immunofluorescent stained tissues. Colorectal liver metastasis microtissues containing up to 30,000 cells were generated and segmented in 5-10 minutes per microtissue [14]. mTmG and Dendra2 fluorescent markers were used to accurately distinguish WT and cancer cells (Fig. 4I). Using a Ki67 staining, NucLogic determined the percentage of Ki67-positive WT and cancer cells, revealing that most cancer cells are in a proliferative state, whereas WT cells largely were not (67% ± 20% versus 10% ± 10%; Fig. 4I-J). Furthermore, NucLogic was used to calculate the average internuclear distance of each nucleus to its three nearest neighbors, using its built-in K-nearest neighbor (KNN) feature. Cancer cells exhibited a lower internuclear distance than WT cells, indicating higher cell density (Fig. 4K). Additionally, non-interacting cancer cells showed significantly greater density than interacting cancer cells, suggesting that cells at the interaction interface may be exposed to different mechanical forces than those further away (Fig. 4K). Together, these analyses demonstrate that NucLogic enables non-experts to perform scalable, single-cell-resolution profiling of both cell-intrinsic and spatial phenotypes, uncovering novel biologically meaningful patterns.

## Discussion

Here, we introduce NucLogic, a generalizable algorithm for 3D nuclear segmentation that uses a logic-based geometric ruleset to stitch 2D masks into accurate 3D volumes. Current state-of-the-art segmentation methods [1,3,8,15] fail to segment nuclei in complex densely packed tissues exhibiting low SNR and z-dependent intensity loss. As a result, robust 3D segmentation has remained poorly suited for routine timelapse and confocal microscopy experiments, restricting the broader adoption of quantitative 3D image analysis [6,7,16,17]. NucLogic addresses this gap by leveraging the robustness of existing 2D segmentation approaches with a novel stitching algorithm to obtain accurate nuclear segmentation, even in such demanding samples. NucLogic first performs rudimentary stitching by linking 2D nuclear segmentations across consecutive slices based on spatial overlap. It then applies a set of human-interpretable geometric rules, based on cross-slice area, intensity, and orientation patterns, to correct z-undersegmentation errors by splitting incorrectly merged nuclei at the appropriate slice. Additionally, a separate correction step resolves 2D undersegmentation, in which touching nuclei sharing a single 2D mask are identified and split at the shared interfacing slice, guided by 3D orientation vectors fitted to each nucleus.

Our approach substantially outperforms existing methods in both accuracy and speed on complex samples. On synthetic data, NucLogic accurately segmented nuclei throughout the entire sample, whereas existing methods failed to detect nuclei in low-SNR regions. This pattern held true for experimental data as well, particularly in low-quality, densely packed tissues, where NucLogic consistently achieves F1 scores above 0.9, driven predominantly by high recall. Moreover, NucLogic segmented large volumes approximately 3-fold faster than competing algorithms. As the imaging field moves toward smart-microscopy, the speed of segmentation algorithms becomes increasingly important to make on-the-fly acquisition decisions [10]. Fast and accurate algorithms such as NucLogic are particularly well-suited for this transition. For example, active cell elimination during cell competition is a rare transient event that is easily missed at screening imaging intervals [18]. By continuously monitoring population-level phenotype ratios, NucLogic could enable a smart microscope setup to detect the onset of decline in the less fit population within each organoid and dynamically increase temporal resolution to capture the elimination process in detail.

The geometric rules described here could be used for the 3D segmentation of any spherical or ellipsoidal structure. Future work could extend on NucLogic’s underlying principles to segment intracellular organelles, such as vesicles [19], lipid droplets [20,21], or biomolecular condensates [22], especially considering these are often acquired under similar challenging conditions characterized by low-SNR and limited z-resolution. The underlying stitching algorithm of NucLogic is inherently size-agnostic and capable of stitching objects of heterogeneous size within a single sample. This depends, however, largely on generating correspondingly heterogeneous 2D segmentations, a task for which Cellpose-SAM is likely not optimally suited. The stitching algorithm itself is agnostic to the underlying 2D segmentation method, and could therefore be paired with simple watershed segmentation approaches or other field-specific tools better suited to a structure’s size range.

Single-cell-resolution analysis of 3D biological images has seen limited adoption across the broader biology field, largely because existing methods remain inaccessible to non-experts [7]. To address this, NucLogic includes a GUI that integrates a wide range of features designed to be accessible for non-expert users. These functionalities enable extensive data mining of complex datasets at single-cell resolution to profile both cell-intrinsic and spatial phenotypes, aiding in the discovery of novel biologically meaningful patterns. For instance, we show that these features can distinguish WT from cancer cells based on fluorophore expression and quantify the proportion of Ki67-postive cells within each population. More broadly, NucLogic can measure additional properties such as the nuclear-to-cytoplasmic intensity ratio of any imaged signal and can integrate features such as local cell density and nuclear morphology, enabling the identification of relevant subpopulations from arbitrary combinations of staining data.

Acquiring accurate 2D segmentation is a prerequisite for our approach, meaning errors introduced at this stage can propagate into the final 3D reconstruction. However, training accurate 2D segmentation models is considerably easier than that required for other existing 3D segmentation approaches, making NucLogic accessible to a broader range of researchers [11,23,24]. A key limitation of NucLogic is that its current geometric rules can fail to produce robust segmentation when z-resolution is very high, as intra-nuclear shape variation may be misinterpreted as inter-nuclear boundaries, resulting in over-segmentation. This limitation was observed in the benchmarking of ultra-dense expansion-microscopy-imaged zebrafish brain sample. However, this can be mitigated by downscaling the image data prior to segmentation, which improves both the segmentation accuracy and computational speed. Alternatively, because NucLogic is built on human-interpretable rules, these rules can be adapted to better match the expected nuclear geometry in such samples. For example, the calculation of *S_break_* (Eq. 1) could be adapted to decrease the weights of the area score to mitigate the effect of intra-nuclear size variations, while still being able to find inter-nuclear shape and orientation changes. This flexibility highlights a key advantage of rule-based approaches over the black-box nature of fully AI-driven 3D segmentation methods [25,26]. Taken together, NucLogic has the potential to make reproducible, single-cell-resolution analysis of complex, low-SNR, multi-dimensional imaging data broadly accessible to biologists, facilitating the discovery of novel spatiotemporal patterns across large-scale datasets.

## Acknowledgements

This work was supported by the European Research Council through Consolidator grant CELL-FIT No. 101169646, Dutch Cancer Society (KWF) No. 2024-3 EXPL / 16529 and Utrecht University focus area Applied Data Science. We thank Valentine Jansen for important discussions on the human-interpretable geometric rules. We thank Rutger Kok and Matthew Smith for valuable discussions and feedback on the manuscript. We thank Elise Mijderwijk, Maeve Troost, Maria Lamprou, and Merel van Luyk for testing the GUI and providing helpful suggestions.

## Methods

### Cellpose-SAM 2D segmentation

Cellpose-SAM models used to obtain 2D segmentations were made by fine-tuning the pretrained generalist Cellpose-SAM checkpoint for 100 epochs with a learning rate between 1e-04 - 1e-05 [1]. Default parameters for batch size and training and validation split were used, determined by the built-in GUI training from Cellpose-4. Training data consisted of up to 25 manually annotated z-slices. Training data was held-out from validation experiments and analysis.

Synthetic and organoid timelapse data were segmented using the “Low quality” model available on GitHub and Zenodo (see data and code availability). Specific models were also trained and made available for the Drosophila embryo, zebrafish embryo, and microtissue (DAPI stained) data. Zebrafish brain data was segmented using Cellpose-SAM’s cpsam_v2 generalist model. These models achieve robust 2D segmentation of nuclear staining or markers across different levels of SNR.

Applying 2D Cellpose-SAM models to slices immediately above or below the foreground can occasionally result in the segmentation of a single, oversized mask that captures the PSF blur surrounding the true signal, rather than an individual nucleus. Such masks are substantially larger than that expected of individual nuclei but are created because Cellpose-SAM is largely size-agnostic. To prevent these artifacts from interfering with 3D reconstruction, all masks exceeding 15 times the median 2D nuclear area within the stack are filtered out.

### Stitching 2D to 3D

2D masks are assembled into 3D masks using a rudimentary stitching approach (Fig. 1A). First, 2D masks larger than 15x the median 2D mask area are discarded, as these are likely 2D segmentation mistakes that arise from faulty background detection in near-empty slices. Next, a KDTree is built of each z-slice for nearest neighbor queries using SciPy [27]. A recursive function uses a bottom-up approach to find the nearest unassigned neighbor within a certain distance threshold (10 pixels by default) on the consecutive slice. When the intersection over union (IoU) of these objects is above 0.3, the masks are assigned the same 3D label, and the next slice is queried until no masks match the description. When a smaller mask is fully contained within a larger mask, its overlap is set to 1. Because only unassigned 2D masks can be considered by the nearest neighbor query, no 2D masks are shared between 3D masks. In this way, all 2D masks are assigned a 3D label.

To limit memory usage and optimize algorithmic speed, all masks data is stored as specialized data classes. This class can store information on 2D information: label, centroid, area, per-channel mean intensity and pixel coordinates, and 3D characteristics: label, centroid, volume, mean intensity, lineage of the parents, break scores, pixel coordinate sets and touching neighbors. Because pixel coordinates are stored only for segmented voxels, as compact integer indices, memory scales with the number of nuclear voxels rather than with the full image volume, which is predominantly background. In addition, the iterative stitching, breaking and merging steps modify these records directly instead of generating a full-size label image at each step. Only after the full pipeline has completed, this representation is rendered into a 16-bit or 32-bit NumPy label image matching the input volume, with each 3D nucleus assigned a unique integer [28].

### Calculating per z-slice break scores and break points

Overall, NucLogic splits densely z-undersegmented nuclei into individual masks by assessing each nucleus against two geometric and intensity-based criteria, which determine whether a split is required and, if so, at which slice it should occur.

To compute the area and intensity-dependent break score, the respective per-slice profiles are inverted, therefore minima become maxima, and valleys can be detected using a peak finding algorithm as implemented by SciPy (Fig. 1B) [27]. Valleys are then normalized against the area of that slice, hence the score reflects relative constriction rather than absolute magnitude to compare nuclei of differing sizes. Because prominence of this type of data is relatively low, both the area and intensity break score are multiplied by 10 before calculating the final break score (*S_break_*). The IoU break score is defined as 1 – IoU, calculated per-slice as mentioned above, to stimulate breaking of poorly overlapping slices. Break scores of the top and bottom slice of a 3D nucleus are always defined as 0, as they have no interface to compare the scores to.

To find geometry related breakpoints, a 3D vector is fitted to the centroids of the three bottom-most slices of each mask using least-squares regression (Fig. 1B). This vector describes the expected trajectory or orientation of the lower nucleus. Next, the distance between each slice centroid and this vector is calculated, and its second derivative is used to identify the rate of change of this distance along z. This rate of change is maximal at the point of greatest mismatch between two z-stacked nuclei, corresponding to the optimal break point. This score is calculated independently in both a bottom-up and top-down direction for every nucleus.

Finally, these individual break scores are combined into a single, final per-slice break score following Equation 1:

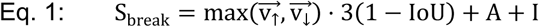

Taking the maximum, rather than combining the bottom-up and top-down scores directly, ensures that each break point is identified independently. When three or more nuclei are stacked, both vectors can register a break signal simultaneously at different points along the stack; summing or multiplying these scores would convolute the two signals into a single, blended value that no longer clearly corresponds to either individual break point. Selecting the maximum instead preserves each break point as a distinct, separately resolvable signal. The vector score is multiplied with the IoU score, as their combination best captures changes in nuclear orientation and trajectory. The area and intensity scores are added rather than multiplied to reflect the hourglass-like pattern. This additive, rather than multiplicative, combination allows each pattern to be detected separately, rather than requiring both to coincide at the same slice.

The *S_break_* of each slice is compared against a set *T_break_*, and a break is made between the slice with the highest *S_break_* exceeding this threshold and its highest-scoring neighboring slice. *T_break_* is derived from *T_original_*, which can be set by the user to fine-tune breaking sensitivity, and the elongation of the mask, following Equation 2.

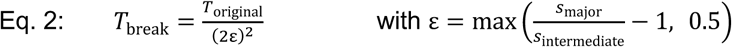

Elongation is calculated using the major and intermediate axes of the mask to specifically capture Z-dependent elongation. Consequently, nuclei that do not represent a sphere-like shape are more prone to breaking than those that remain approximately spherical. All nuclei are iteratively checked, broken, and reanalyzed until there are no remaining nuclei that contain slices exceeding *T_break_*.

### Splitting shared 2D segmentation to prevent undersegmentation

2D segmentation of z slices that contain two neighboring nuclei can result in undersegmentation. To address this, NucLogic finds, for each nucleus, all touching neighbors in a 3D volume. For each touching pair, a 3D line is fitted through the slice centroids of both nuclei, excluding the touching interface slice in question. These vectors are used to predict two centroid positions in the interfacing slice, under the assumption that the interfacing slice is undersegmented. The interfacing slice is then split along a line perpendicular to the vector connecting the two predicted centroids. The two resulting 2D masks both give a new centroid position. Equation 3 is applied to check whether this split improves segmentation accuracy.

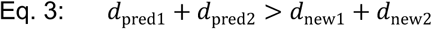

*d_pred1_* and *d_pred2_* represent the distance between the original centroid and the newly predicted centroids. *d_new1_* and *d_new2_* are the distances between the newly predicted centroids and the newly created centroids. Consequently, a split is only accepted and retained when the newly predicted centroids better represent the expected nuclear trajectory. This procedure is applied to the interfacing slices of both nuclei, and in case the first layer is split it is repeated on the second neighboring slice.

### Filtering of 3D masks

Following stitching and splitting, the 3D attributes of each mask are recalculated and stored within the specialized 3D data structure. A final cleanup step is then performed to remove small background artifacts and correct potentially over-segmented nuclei. To this end, all masks comprising only one or two z-slices are compared against neighboring nuclei in adjacent slices for potential IoU overlap, using similar approaches as described above. Small masks with more than 0.7 IoU relative to a neighboring nucleus above or below are merged with that nucleus, while those that do not meet this criterion are discarded as likely background artifacts. The final generated 3D data class is then transformed into a uint16 or uint32 NumPy array, storing up to 65,536 or 4,294,967,295 individually segmented nuclei, respectively.

### Automatic 3D timelapse cropping

To reduce computing time and enable isolation of a single 3D structure (e.g., an organoid) from surrounding objects in the same field of view, NucLogic includes an automated method for cropping 3D imaging data. For each timelapse, the organoid is first cropped in XY using maximum z-projections of the nuclear marker channel(s). Organoids of interest are identified in the initial frame using a combination of the Gaussian blur and triangle thresholding methods implemented by scikit-image [29,30]. The resulting mask is dilated and filled to identify connected components using OpenCV [31]. The object closest to the image center is automatically selected as the seed object for tracking. The seed mask initializes Meta’s SAM 2 video segmentation model, which propagates the organoid masks across all timepoints [12]. To correct tracking failures — which can arise from dimming regions caused by organoid growth or changes in sample depth along z — each frame is monitored for sudden drops in mask area (>30% relative to the preceding frame). When detected, the model’s own mask from the last successfully tracked frame is used to re-seed propagation from that point forwards. The resulting masks are dilated by a set number of pixels to ensure complete inclusion of the organoids. Alternatively, when the SAM model cannot infer correct XY cropping of non-timelapse data, a manual XY cropping mode can be used to draw a region of interest on the sample.

Next, each frame is also cropped in z by computing Moran’s I for spatial autocorrelation on each row of the XZ maximum-intensity projection [32]. In this way, slices that contain background (i.e. no spatial correlation) can be separated from foreground slices. Rows with a Moran’s I above 40% of the frame’s maximum value are marked, and the longest contiguous block of marked rows (allowing gaps up to 6 slices) is selected as the region of interest. A 2-slice margin is added on each side to ensure complete inclusion of the organoid. A row-wise 2D approach on the XZ projection is used, rather than computing Moran’s I in full 3D, to optimize computational speed. Each frame is cropped and padded to a common, minimal shape to construct the final 3D timelapse array.

### Phenotype calling

To determine whether a nucleus belongs to phenotype A or B, the log10 ratio of mean intensities of channels A and B is calculated to approximate relative intensity levels for each nucleus. In this way, each quantification is normalized against z-dependent intensity drops. A bimodal distribution is expected in the histogram of the log10 intensity ratios, reflecting the two underlying phenotype populations. A two-component Gaussian mixture model, implemented with scikit-learn, is used to model the two underlying Gaussian distributions [33]. The quadratic formula is then applied to solve for the intersection point of the two fitted Gaussian distributions, which defines the automatic cutoff between populations. For samples where one phenotype is substantially more abundant than the other (standard deviation of log-ratios ≤ 0.55), the cutoff is instead set just beyond the observed data range, effectively assigning all nuclei to the dominant phenotype. As an additional safeguard, if more than 97% of nuclei are classified into a single phenotype under the automatic cutoff, all nuclei are assigned to that phenotype to avoid misclassification driven by noise.

Alternatively, the NucLogic UI also supports setting a custom cutoff to identify populations that cannot be captured by a bimodal distribution, for example when one population is scarce. To facilitate determination of the correct threshold, the sample can be opened in napari directly from the NucLogic UI. Here, a custom plugin loads NucLogic-generated statistics and provides an interactive slider to manually identify the appropriate cutoff for that population.

### Cell-intrinsic and spatial statistics

A wide variety of cell-intrinsic and spatial statistics can be computed per nucleus using the NucLogic UI. Cell-intrinsic properties, such as maximum intensity, elongation and volume, are measured using scikit-image regionprops [29]. All size-dependent properties are calculated using voxel size metadata extracted from the input file, or a user-defined voxel size when such metadata is unavailable. These metrics can be calculated from the nuclear mask and can also be estimated for the cytoplasm or whole-cell mask. Whole-cell masks are generated by a Euclidean distance transform (EDT) of the background with the nuclear masks as seeds [27]. The voxel size is used as a sampling rate to ensure correct handling of anisotropic imaging volumes. Every background voxel within a user-defined distance (in µm) of a nucleus is assigned the label of the nearest nuclear voxel, hence adjacent nuclei compete for the intervening space and expanded masks never overlap. The cytoplasmic mask is generated by removing the nuclear mask pixels from the whole-cell mask, forming a shell around each nucleus. Expanded masks can be stored for later visualization.

Spatial properties, such as cell-cell interaction identification and cell density, can be calculated in two ways; (1) A KNN approach using a kd-tree on 3D centroid positions, or (2) by identifying pairs of touching cells from the EDT-dilated masks [27,33]. The phenotype similarity score is calculated as the fraction of a cell’s neighbors that share its phenotype, ranging from 0 when no neighbor matches to 1 when all do. The identity of all neighbors is stored, enabling additional custom downstream analysis outside the NucLogic UI.

### Data visualization and export

Image data and segmentations can be inspected directly from the NucLogic UI, which opens selected samples in a napari viewer [34]. Each imaging channel is added as a separate image layer with a specified color map, and contrast limits are set automatically. Segmentation masks are rendered as smooth surfaces, and any whole-cell, cytoplasmic, or phenotype-specific masks can be added. Each layer is scaled by the voxel size retrieved from the image metadata for correct viewing of anisotropic volumes. The napari-animation plugin is included in the NucLogic environment to enable generation of animations of the imaging data and segmentations.

Within the viewer, a custom napari plugin links each label layer to that sample’s statistics table. Any property can be selected, upon which a histogram and a range slider are shown, and the displayed nuclei or cells are restricted in place to those falling inside every active range. This allows populations to be identified visually and interactively. The resulting selection can be split off into a new layer, saved as a compressed TIFF carrying the same voxel size and time metadata as the rest of the pipeline, or annotated back into the statistics table as an additional column for downstream use.

All properties are written per sample as tab-separated (TSV) files. Integer nuclear labels corresponding to the segmentation masks are stored automatically, along with centroid position, volume, and mean intensity for each imaging channel. Additionally, the export data tab can be used to concatenate the data of specified samples into a single TSV. Alternatively, a summary table can be generated, reporting the number of cells assigned to each phenotype per timepoint.

### Benchmarking against existing segmentation methods

NucLogic was benchmarked against Cellpose-SAM [1,2], StarDist [3,4], and u-Segment3D [8]. Cellpose-SAM was applied in 3D mode and run using a flow3D_smooth value of 0.4, and both normalization and resample set to True, as these parameters produced the most accurate segmentation. Models were trained as described above and the best-performing model was used for benchmarking of each sample. For StarDist segmentation, the pretrained 3D segmentation model was used. Each image was normalized, and model prediction was performed using n_tiles determined by StarDist’s guess_n_tiles function. For u-Segment3D, we tested both the indirect and direct methods using both Cellpose 3 and Cellpose-SAM as a backbone. The data reported here were generated using the indirect method with Cellpose-SAM, using the pretrained models described above, as this configuration produced the most accurate results. For this we used the following parameters on the XY presegmented volume: dtform_method = edt, binary_closing = 0, do_mp = False, thresh_factor = 1, and smooth_sigma = 0.

To compare all segmentation methods, we calculated the F1, recall, and precision scores based on centroid accuracy. From the ground truth segmentations, each centroid position is compared against the test mask. A non-zero label in the test mask that has not yet been claimed counts as a true positive, a label already assigned to an earlier ground-truth nucleus counts as a false positive (over segmentation), and a background voxel in the test mask counts as a false negative, while any predicted object never hit by a ground-truth centroid counts as a false positive. Additionally, F1, recall, and precision scores were also calculated using various IoU thresholds following the average_precision metric as provided by Cellpose following the Hungarian algorithm (Fig. S2 and S4) [2].

Benchmarking was performed on a standard workstation computer using an Intel Core i7-14700 CPU, 32 GB of DDR5 memory, and an NVIDIA RTX 5070 Ti GPU.

### Synthetic data generation

To generate synthetic organoids, ellipsoidal nuclei of a random z, y, and x size (mean radius of 5 × 12 × 12 voxels, respectively) were generated. 300 of these cells were placed on the shell of a 380-voxel diameter sphere with their flat side facing the center of the sphere to resemble cystic organoids. A minimum gap of 4 voxels between each nucleus was maintained. The volume was then rescaled to voxel sizes representative of most confocal organoid timelapse imaging (2.5 × 0.64 × 0.64 μm per voxel). Each nucleus was assigned a mean intensity drawn from a normal distribution centered at 150 (SD 30), with per-voxel variation of SD 5. To mimic PSF spread and tissue scattering, an anisotropic Gaussian whose width increases with depth was applied following Equation 4, with *d* as the normalized z-depth.

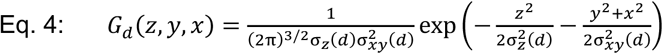

A depth-dependent signal loss was applied following a square root decay modulated by a logistic term, using equation 5.

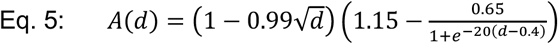

We set the midpoint of the logistic term at *d* = 0.4 and its steepness to 20, so that the deeper half of the synthetic organoids were strongly degraded, as this most closely matched our real-world data. At last, random poison shot noise, Gaussian read noise, and a constant camera offset were added.

### Expansion-assisted imaging and DAPI staining

Wild-type zebrafish (*Danio rerio*) were housed at the zebrafish facility of Utrecht University under a 14 h light/10 h dark cycle at 28 °C. Embryos were collected and kept at the same conditions until 5 days post-fertilization (dpf). Then they were anesthetized in ice-cold water and fixed overnight in 4% paraformaldehyde (PFA) in 1× phosphate-buffered saline (PBS) and stored in 70% ethanol.

Expansion microscopy was performed based on the protocol described by Close et al. [35], using the gel solution 1 described by Ruetten et al. [36], with the modifications described below. Following fixation, larvae were depigmented using 3% H₂O₂ in 0.8% KOH for a maximum of 5 min, as described previously [37]. The depigmented whole larvae were subsequently mounted and equilibrated in MOPS– Triton buffer (0.5% Triton-X100, pH 7.8) before anchoring. For anchoring, specimens were incubated in 1 mg mL⁻¹ melphalan-AcX supplemented with 0.1 mg mL⁻¹ additional AcX, freshly diluted in MOPS– Triton buffer. The gel solution was prepared according to Ruetten et al. [36], with 1% bis-acrylamide. Before gelation, specimens were preincubated in gel solution for three consecutive 20-min incubations at 4 °C. Gelation was subsequently performed at 37 °C for 1.5 hours. Specimens were digested at 50 °C overnight, followed by an additional 4-h incubation in proteinase K (ProK) buffer containing 0.3% SDS. Gels were subsequently equilibrated in 1× PBS for 2x expansion.

For nuclear staining, gels were incubated with DAPI (1 mg mL⁻¹ stock solution in H₂O; Sigma-Aldrich, D9542-1MG; stored at 4 °C) diluted 1:1000 in 1× PBS at 4 °C over the weekend. Before imaging, specimens were equilibrated to room temperature for at least 1 h in fresh 1× PBS. Residual DAPI was removed by washing the specimens three times for 10 min in fresh 1× PBS at room temperature.

### Zebrafish brain imaging

Samples were mounted in 35-mm glass-bottom imaging dishes (ibidi) coated with poly-L-lysine and imaged using a spinning-disk confocal microscope consisting of an Eclipse Ti2-E microscope with Perfect Focus System (PFS; Nikon, Japan) and a CSU-W1-T1 spinning-disk confocal scanner (Yokogawa, Japan), equipped with a 40× water-immersion objective (NA 1.15; Nikon). Pixel size is 0.167 µm.

To obtain a dense and homogeneous population of DAPI-positive nuclei, z-stacks were acquired from the border region between the midbrain and hindbrain using a 405-nm Vortran Stradus laser (100 mW; Vortran, USA) and the corresponding DAPI filter set (ET460/50m). Images were acquired with a z-step of 1 µm.

### Microtissue culture and imaging

Mixed liver metastasis microtissues were generated by mixing differentiated hepatocyte-like organoids with colorectal cancer organoids according to our previously described protocol [14]. Staining was performed as previously described, using a blocking and antibody buffer of 0.5% triton and 5% BSA in PBS, and a washing buffer of 0.1% Triton in PBS. Ki67 was stained using anti-Ki-67 (1:1000, CAT# 14-5698-82, eBioscience) and Donkey anti-Rat Alexa Fluor Plus 647 1:500, (CAT# A-48272, Thermo Fisher Scientific). Nuclei were labeled using DAPI (1:1000, CAT# D416050, Toronto Research Chemicals). Samples were washed at the end with PBS and put into RapiClear 1.49 clearing solution (Sunjin Labs CAT# RC149001).

Microtissue images were acquired using a Andor BC43 confocal benchtop microscope with Borealis™-illuminated multipoint confocal 405/488/561/638 nm excitation lasers, equipped with a Nikon 20X CFI Plan Apo LD dry immersion objective. Images were taken with a voxel size of 2.5 × 0.64 × 0.64 μm.

### Software design and data availability

The source code of NucLogic and its installation instructions are available at https://github.com/SuijkerbuijkLab/NucLogic and the version used to generate this manuscript is permanently archived on Zenodo (DOI: 10.5281/zenodo.23110802). Code used to generate the analysis and figures of here is included in the same repositories. Additionally, the pretrained Cellpose-SAM models used in this study are available on Zenodo (DOI: 10.5281/zenodo.23110966).

NucLogic supports a broad range of features, including but not limited to those showcased in this paper: image and timelapse cropping, segmentation, phenotype calling, cell-intrinsic and spatial statistics calculation, PSS-based interaction scoring using EDT-based dilation or K-nearest-neighbor analysis, data visualization, and data export. All functionalities are accessible through the GUI to non-experts, requiring no coding experience. Data visualization is handled through built-in napari [34]. A custom napari-plugin is included, enabling visualization and filtering of segmented nuclei based on any NucLogic-generated statistic. Importantly, NucLogic features automatic update detection and installation, ensuring users have immediate access to future feature releases.

Software is written in Python 3.11, using Pixi environments for cross-platform installation and PySide6 for the GUI, and uses established software packages: PyTorch v2.10 with CUDA 12.8 [38], LibPySal [39], scikit-image [29], scikit-learn [33], OpenCV [31], NumPy [28], SciPy [27], pandas [40], Matplotlib [41], Seaborn [42], napari [34], Cellpose-SAM [1], and SAM 2 [12]. Due to the PyTorch and CUDA version, NucLogic supports most modern workstations equipped with an NVIDIA GPU produced from 2019 onwards. Alternatively, NucLogic can be installed on Windows or Linux computers without a GPU to calculate statistics or visualize previously segmented samples (segmentation itself is extremely slow without GPU acceleration).

The GUI supports interactive sample selection and accepts most major imaging file formats (tiff, zarr, ims, nd2, czi, lif, etc.) as input. Voxel size metadata is automatically read to accurately calculate size-dependent statistics. Segmentations are saved in TIFF format, and statistical output is saved as TSV files. Users select which statistics to calculate directly through the GUI, and both full and summarized datasets can be exported Drosophila and zebrafish embryo data was obtained from [11]. The organoid, zebrafish brain, and microtissue data reported in this paper are available upon request from the lead author.

## Supplemental materials

**Figure S1:**
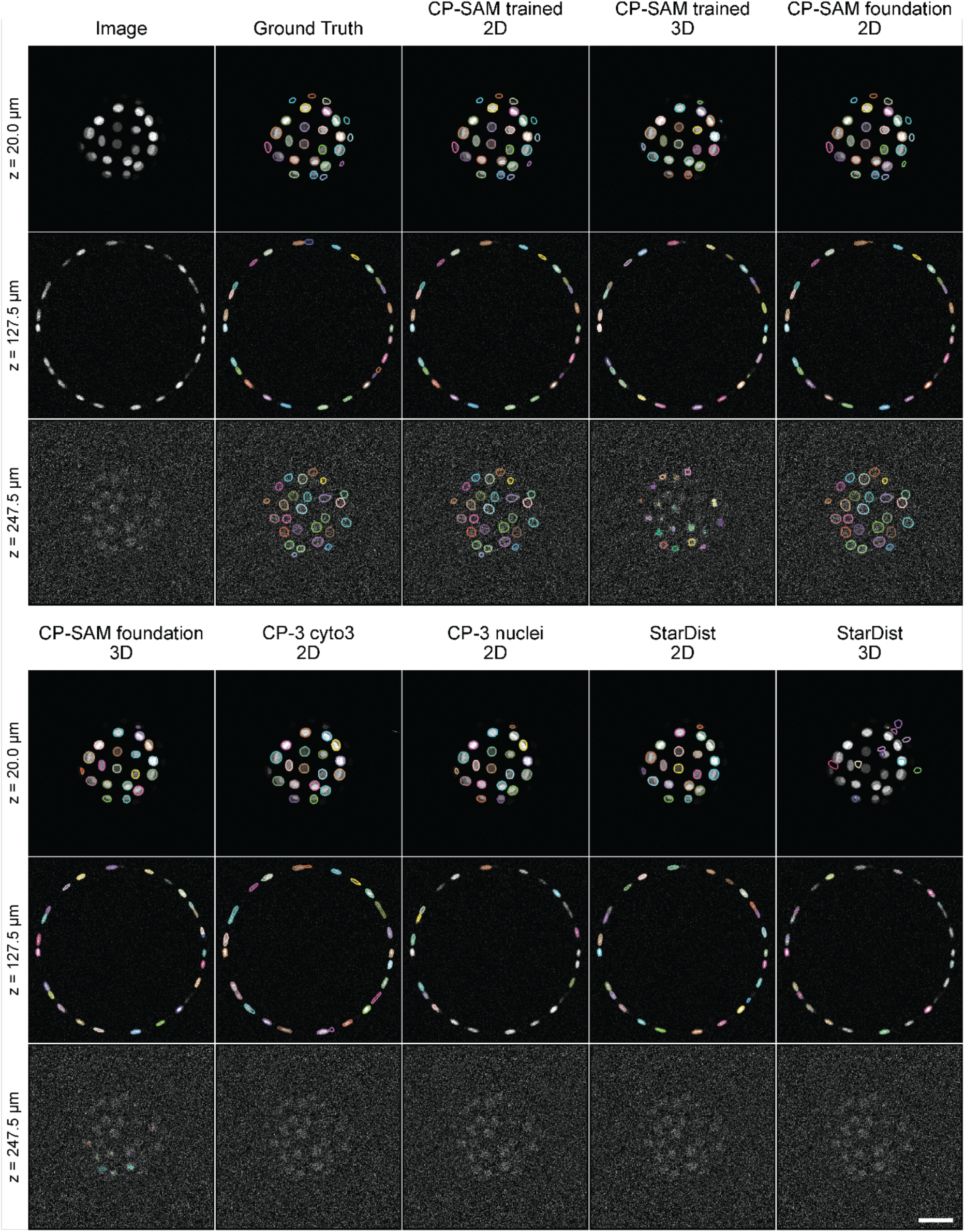
Comparison of 2D segmentation methods across z-dependent SNR variation. Raw image data are shown alongside ground-truth contours and segmentations from: Cellpose-SAM in 2D and 3D mode (fine-tuned trained model and foundation model), Cellpose-3 using the cyto3 and nuclei models in 2D mode, and StarDist in 2D and 3D mode. Three representative slices were selected at different z-depths to compare segmentation performance across varying SNR levels. Scale bar = 50 µm.

**Figure S2:**
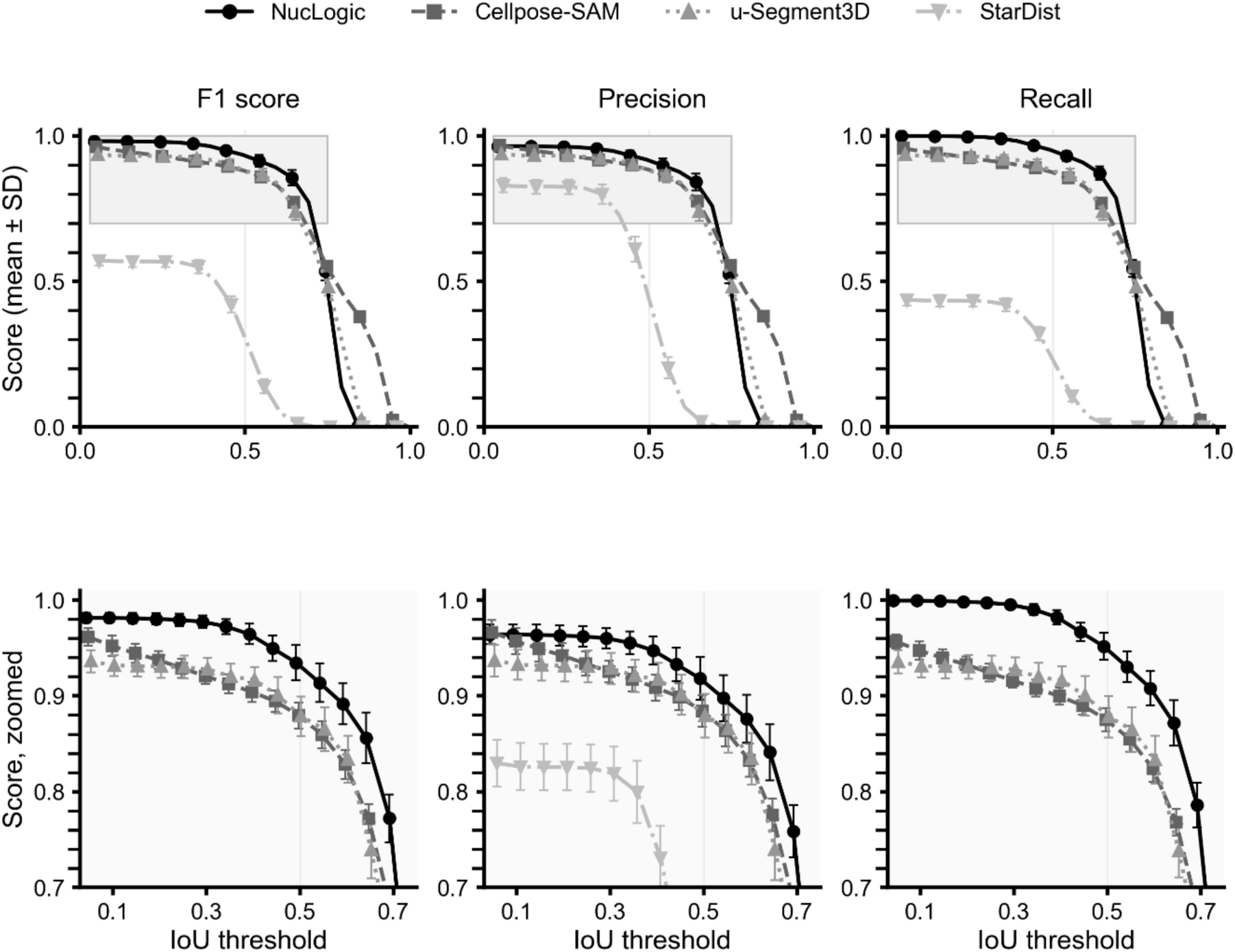
Segmentation accuracy of synthetic data. Synthetic organoids were segmented using NucLogic, Cellpose-SAM, u-Segment3D, and StarDist. F1, precision, and recall scores were measured using the Hungarian algorithm for increasing IoU thresholds. Top panel shows plots of the full data overview. Bottom panel shows zoomed graphs, data represented as mean ± SD (N = 20).

**Figure S3:**
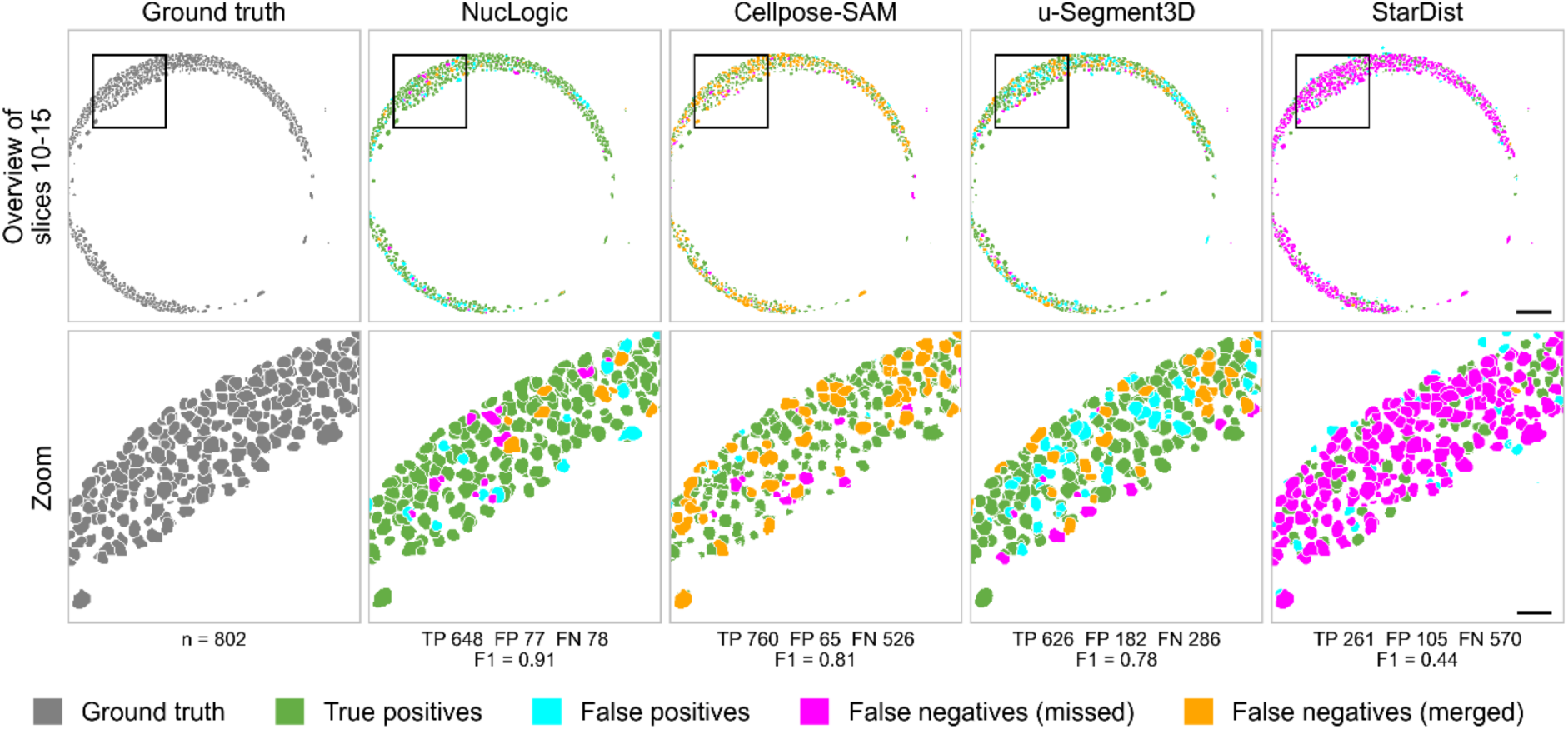
Segmentation accuracy of low SNR slices in zebrafish embryo. Ground truth, NucLogic, Cellpose-SAM, u-Segment3D and StarDist segmentations of the zebrafish embryo from the NIS3D dataset. Top: maximum z-projection of slices 10–15 (15 µm, 2.5 µm z-step). These slices are far from the objective and therefore have a low signal-to-noise ratio (SNR). True positive (TP, green), false positives (FP, cyan). Ground-truth nuclei whose centroid falls on background are missed false negatives (FN, magenta). Merged false negatives (orange) are ground-truth nuclei whose centroid falls inside a predicted nucleus that has already been matched to another ground-truth nucleus, meaning several nuclei were segmented as one object. Bottom: zoom of the respective segmentations. The numbers of TP, FP and FN and the F1 score are shown beneath each panel. They are calculated over the full field of view for all nuclei with at least one voxel in slices 10–15. Scale bars: 100 µm (overview) and 25 µm (zoom).

**Figure S4:**
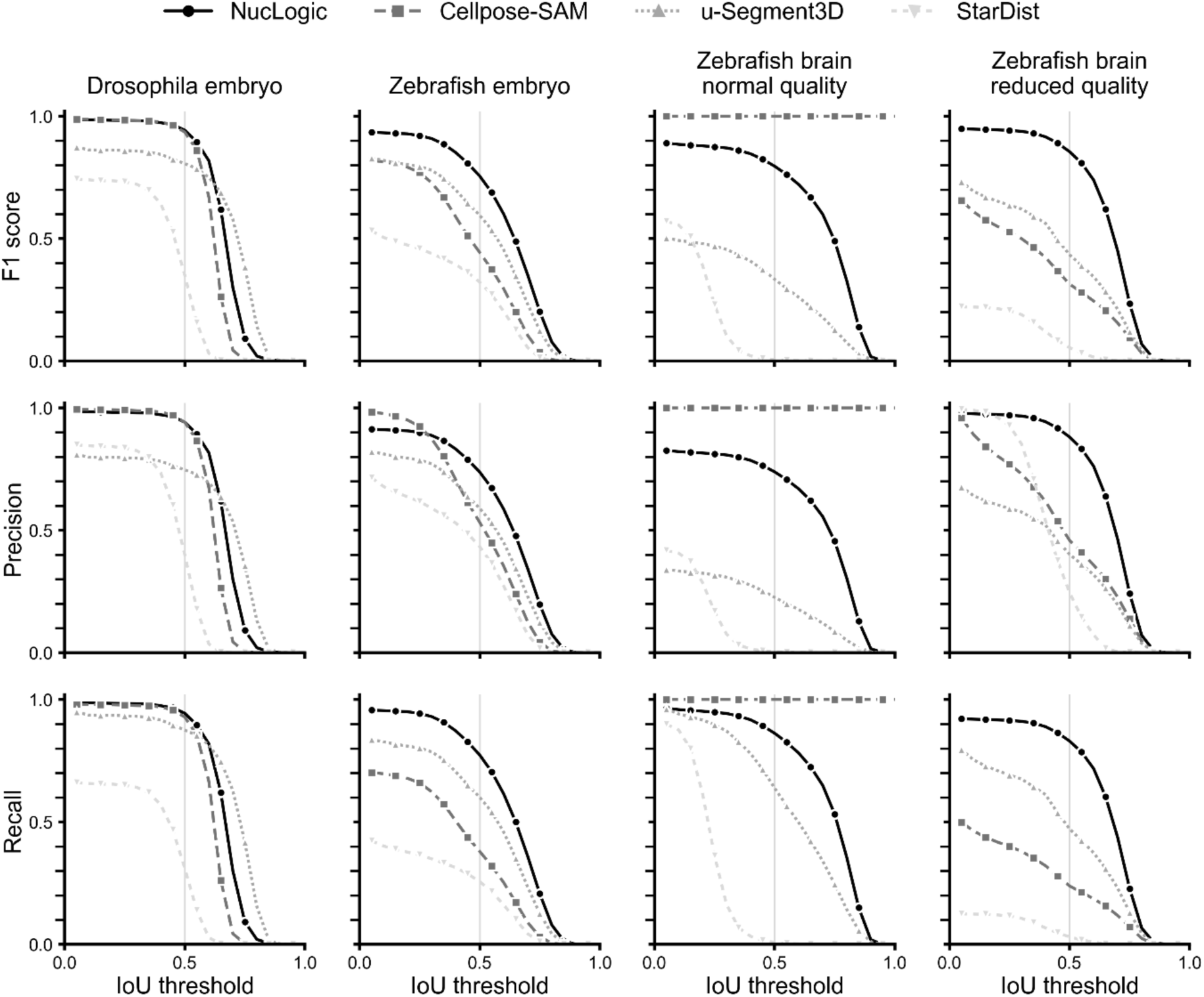
Segmentation accuracy of real-world benchmark data. All samples represented in Fig. 2 (Drosophila embryo, zebrafish embryo, and the zebrafish brain at normal and reduced quality) were segmented using NucLogic, Cellpose-SAM, u-Segment3D, and StarDist. F1 (top), precision (middle), and recall (bottom) scores were measured using the Hungarian algorithm for increasing IoU thresholds.

**Figure S5:**
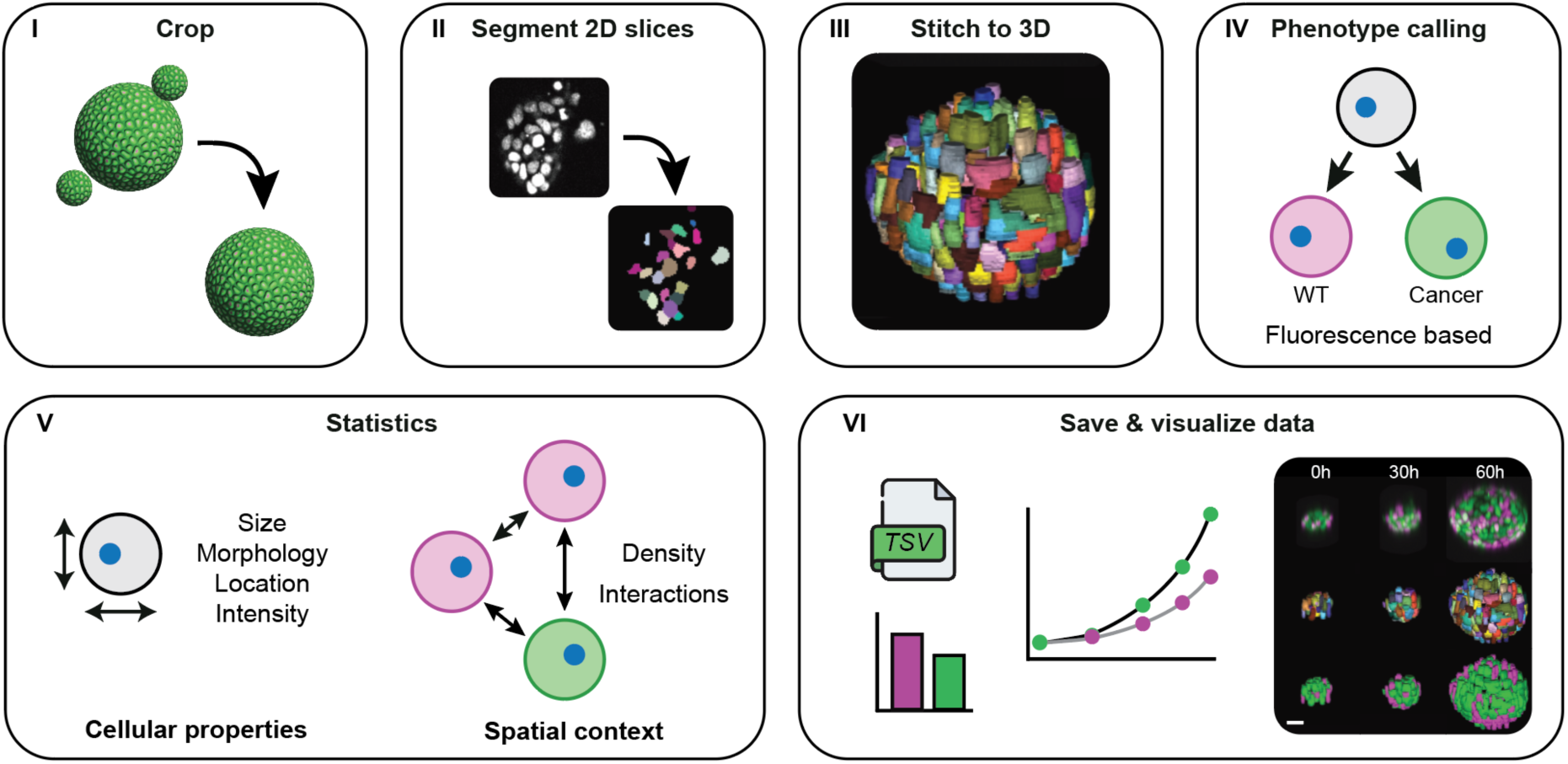
NucLogic complete pipeline overview. The NucLogic GUI enables end-to-end analysis of 3D fixed and timelapse data. (I) Processing begins with cropping of the input data in 3D. (II) The sample is segmented in 2D using Cellpose-SAM and then (III) transformed into 3D masks using NucLogic stitching algorithm. (IV) NucLogic can apply a built-in phenotype-calling algorithm that assigns a phenotype to each segmented nuclei based on any imaged fluorescent marker. (V) A wide range of statistics, including nucleus-intrinsic properties including size, location, morphology, and intensity, as well as spatial context-dependent statistics such as cell density and interactions between neighboring cells can be measured using NucLogic. (VI) The resulting segmentations and their accompanying statistics can be directly visualized by launching napari from the NucLogic UI.

**Figure S6:**
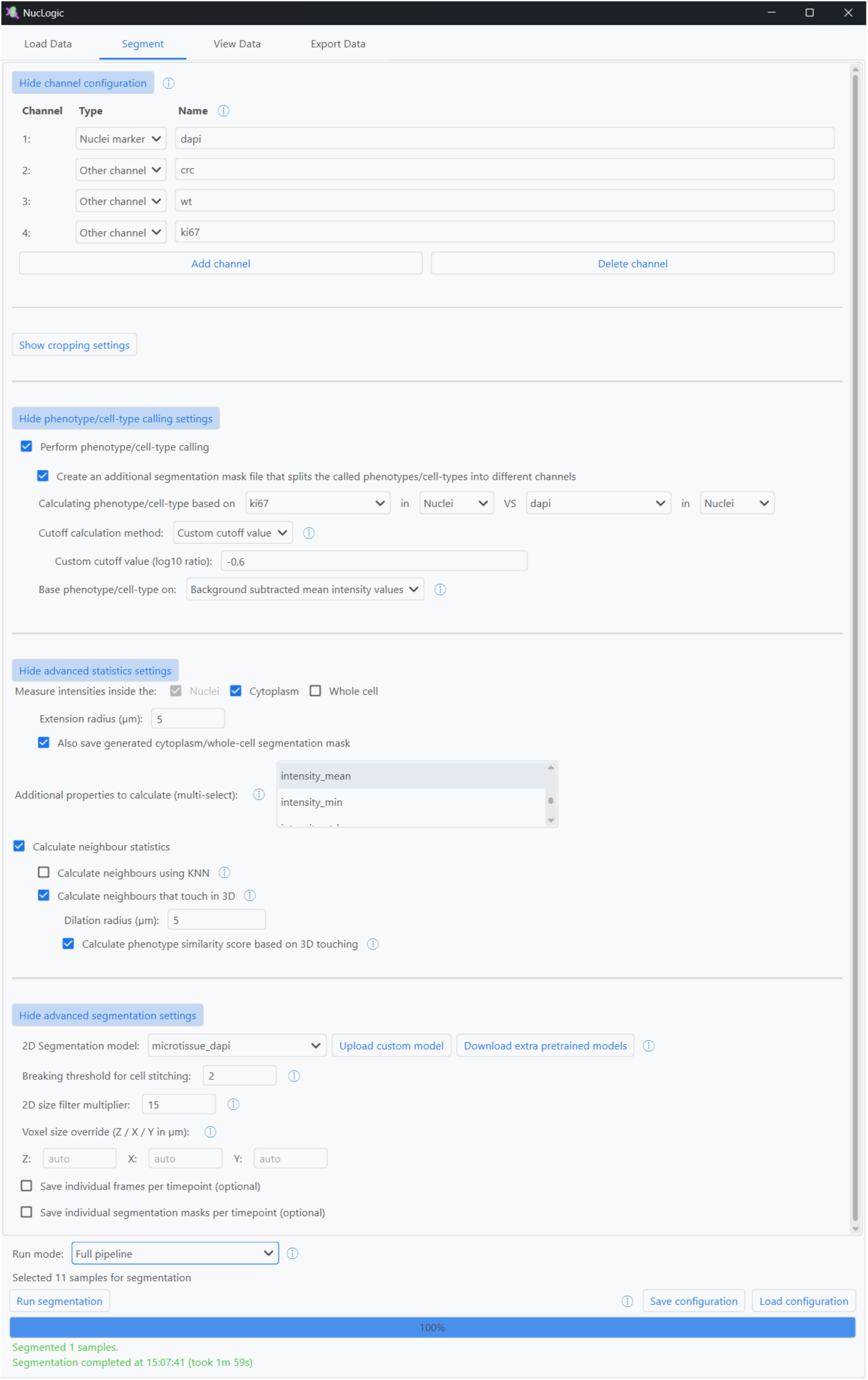
Segmentation and processing tab of the GUI. The segment tab forms the main page of the GUI. The channels of the images can be set, specifying which channel(s) are nuclear markers for segmentation, and which channels are for measuring intensity data. Channel names can be set for easier downstream processing. Cropping settings can be altered to facilitate automatic 3D cropping of timelapse and fixed data, or manual cropping of fixed data. Phenotype calling settings can be set to subdivide segmented cells into different categories based on nuclear, cytoplasmic or whole-cell intensities of all measured channels. This can be done either automatically or with custom thresholds. Either raw or background subtracted values can be used for these measurements. The advanced statistics section is used to get more information on nuclear, cytoplasmic, or whole-cell labels, such as max intensities or shape parameters (elongation, solidity, etc.). Spatial statistics such as neighbors and the phenotype similarity score (PSS) can be found using the KNN or EDT method. Advanced segmentation parameters, such as the 2D Cellpose-SAM model, breaking threshold, and 2D size filter can be set to the needs of the sample. Custom voxel sizes can be set when input images do not contain proper metadata. The run mode can be selected to differentiate what parts of the full NucLogic pipeline need to be performed, such that reclassifying statistics is not dependent of resegmenting the sample. All settings (configuration) can be saved and loaded to guarantee consistency throughout experiments or analysis runs. A progress bar visualizes the progression of the analysis of all selected samples.

**Figure S7:**
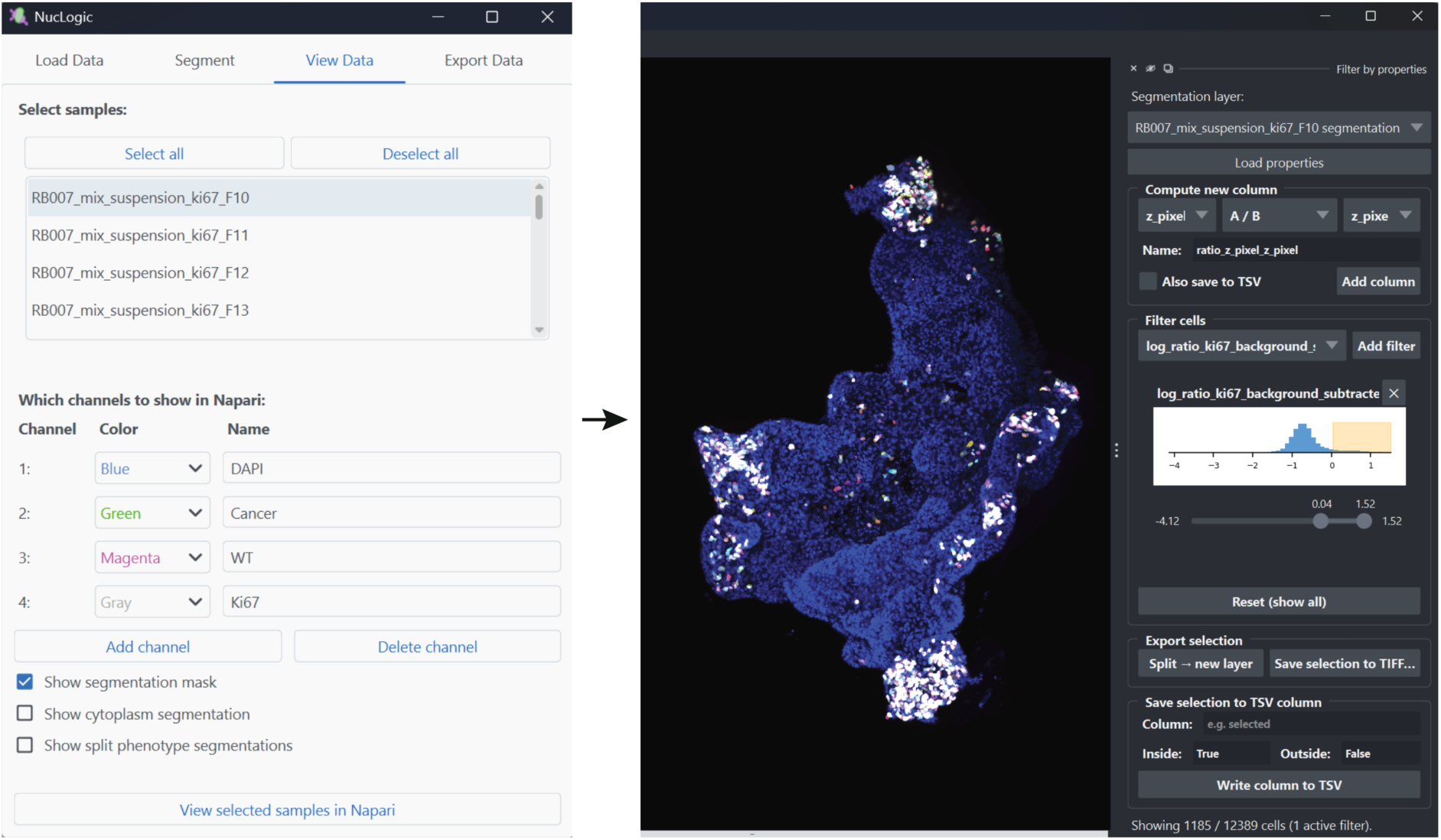
View data tab of the GUI and napari plugin to find phenotype calling thresholds. Left: The view data tab is used to select samples for visualization. The channel colors and names can be set. Nuclear, cytoplasmic, and whole-cell segmentation masks are automatically registered and opened in napari. Any phenotypes that have been called can be visualized as integer masks and as specified colored mask. Right: NucLogic ships with a custom napari plugin that supports finding specific populations of segmented cells. All NucLogic generated statistics can be used as filters. A histogram of the specified statistics is visualized, and a slider is used to select the desired population of cells. Resulting masks can be saved as .TIFF files and newly generated categories can be saved to the TSV files generated by NucLogic.

**Table S1:** Segmentation scores of synthetic organoids represented in Figure 1G.

| **Method** | **F1** | **Precision** | **Recall** | **Segmentation time (s)** |
| --- | --- | --- | --- | --- |
| *NucLogic* | 0.979 ± 0.007 | 0.962 ± 0.011 | 0.997 ± 0.004 | 33.764 ± 1.005 |
| *Cellpose-SAM* | 0.964 ± 0.008 | 0.969 ± 0.012 | 0.959 ± 0.008 | 106.433 ± 1.268 |
| *u-Segment3D* | 0.929 ± 0.012 | 0.930 ± 0.017 | 0.928 ± 0.013 | 47.242 ± 1.420 |
| *StarDist* | 0.568 ± 0.019 | 0.824 ± 0.025 | 0.433 ± 0.021 | 9.997 ± 0.470 |

**Table S2:**
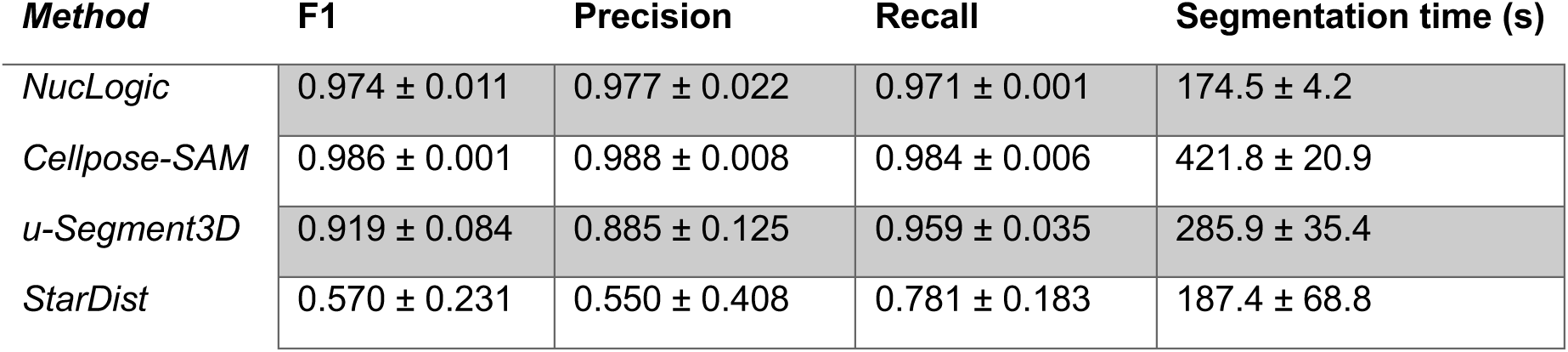
Segmentation scores of NIS3D Drosophila embryos represented in Figure 2A.

| **Method** | **F1** | **Precision** | **Recall** | **Segmentation time (s)** |
| --- | --- | --- | --- | --- |
| *NucLogic* | 0.974 ± 0.011 | 0.977 ± 0.022 | 0.971 ± 0.001 | 174.5 ± 4.2 |
| *Cellpose-SAM* | 0.986 ± 0.001 | 0.988 ± 0.008 | 0.984 ± 0.006 | 421.8 ± 20.9 |
| *u-Segment3D* | 0.919 ± 0.084 | 0.885 ± 0.125 | 0.959 ± 0.035 | 285.9 ± 35.4 |
| *StarDist* | 0.570 ± 0.231 | 0.550 ± 0.408 | 0.781 ± 0.183 | 187.4 ± 68.8 |

**Table S3:**
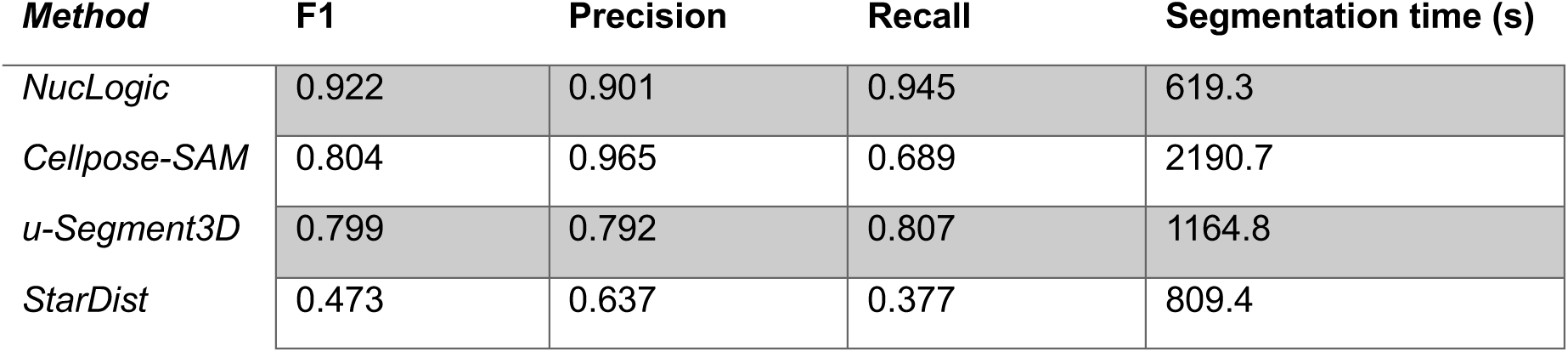
Segmentation scores of NIS3D zebrafish embryo represented in Figure 2B.

| **Method** | **F1** | **Precision** | **Recall** | **Segmentation time (s)** |
| --- | --- | --- | --- | --- |
| *NucLogic* | 0.922 | 0.901 | 0.945 | 619.3 |
| *Cellpose-SAM* | 0.804 | 0.965 | 0.689 | 2190.7 |
| *u-Segment3D* | 0.799 | 0.792 | 0.807 | 1164.8 |
| *StarDist* | 0.473 | 0.637 | 0.377 | 809.4 |

**Table S4:** Segmentation scores of expanded zebrafish brain at normal quality represented in Figure 2C.

| **Method** | **F1** | **Precision** | **Recall** | **Segmentation time (s)** |
| --- | --- | --- | --- | --- |
| *NucLogic* | 0.887 | 0.824 | 0.960 | 424.2 |
| *Cellpose-SAM* | 1 | 1 | 1 | 1725.9 |
| *u-Segment3D* | 0.492 | 0.335 | 0.927 | 4399.2 |
| *StarDist* | 0.378 | 0.274 | 0.611 | 2824.0 |

**Table S5:** Segmentation scores of expanded zebrafish brain at reduced quality represented in Figure 2C.

| **Method** | **F1** | **Precision** | **Recall** | **Segmentation time (s)** |
| --- | --- | --- | --- | --- |
| *NucLogic* | 0.946 | 0.967 | 0.925 | 54.3 |
| *Cellpose-SAM* | 0.623 | 0.891 | 0.479 | 293.3 |
| *u-Segment3D* | 0.644 | 0.598 | 0.697 | 364.1 |
| *StarDist* | 0.189 | 0.972 | 0.105 | 58.4 |

